# Significant East Asian affinity of Chinese Hui genomic structure suggesting their predominant cultural diffusion model in the genetic formation process

**DOI:** 10.1101/2021.01.12.426452

**Authors:** Yan Liu, Junbao Yang, Yingxiang Li, Renkuan Tang, Didi Yuan, Yicheng Wang, Peixin Wang, Shudan Deng, Simei Zeng, Hongliang Li, Gang Chen, Xing Zou, Mengge Wang, Guanglin He

## Abstract

Ancestral origin and genomic history of Chinese Hui people remain to be explored due to the paucity of genome-wide data. Some evidence argued that an eastward migration of Central Asian given rise to modern Hui people, which was inferred as the *demic diffusion hypothesis*, and others favored the *cultural diffusion hypothesis* that posited indigenous East Asian adopted Muslim-like culture and formed the modern culturally different populations. However, the extent to which the observed Hui’s genetic structure was mediated by the movement of people or the assimilation of Muslim culture also remains one of the most contentious puzzles. Analyses of over 700K SNPs in 109 western Chinese individuals (49 Sichuan Hui and 60 geographically close Nanchong Han) together with the available ancient and modern Eurasians allowed us to fully explore the genomic makeup and origin of Huis and neighboring Hans. The results of the traditional and formal admixture-statistics (PCA, ADMIXTURE, and allele-sharing-based *f*-statistics) illuminated a strong genomic affinity between Sichuan Hui and Neolithic-to-modern Northern East Asians, which suggested massive gene influx from East Asian into Sichuan Hui people. Three-way admixture models in the *qpWave/qpAdm* analyses further revealed a small stream of gene influx from western Eurasian related to French or Andronovo into these Hui people, which was further directly confirmed via the admixture event from the temporally different western sources to Hui people in the *qpGraph*-based phylogenetic model, suggesting the key role of cultural diffusion model in the genetic formation of the modern East Asian Hui. ALDER-based admixture date estimation showed that this observed western Eurasian admixture signal was introduced into East Asian Hui during the historic periods, concordant with the extensive western-eastern communication in the Silk Road and historically documented Hui’s migration history. Summarily, although significant cultural differentiation among Hui and their neighbors existed, our genomic analysis showed their strong affinity with modern and ancient Northern East Asians. Our results supported that modern Chinese Hui arose from the mixture of minor western Eurasian ancestry and predominantly East Asian ancestry.

## Introduction

Archeologically and historically attested transcontinental exchange during the past five thousand years in North China and Northwest China played an important role in the formation of modern observed genetic/linguistic/cultural diversity in East Asian people (Dong et al., 2020). The archaeological record showed that wheat/barley agriculture and plant/animal domestication technologies from the Fertile Crescent have spread into North China (Leipe et al., 2019;Dong et al., 2020). Besides, Northern Chinese millet farming technology also has been disseminated into western Eurasia (Leipe et al., 2019;Dong et al., 2020). Zoo-archeological/Phyto-archeological findings and the existing western Bronze packages (metallurgy and chariots, horse, cattle and sheep/goat) in China suggested that the massive western cultural elements were introduced into China in response to the transformation of human subsistence patterns or climate changes (Leipe et al., 2019). Late-Paleocene and early-Holocene genomes from Siberia have revealed several genetically diverse Paleolithic lineages (Ancient Northern Eurasian Mal’ta1 lineage, Ancient North Siberian Yana lineage et al.) and a typical West-to-East genetic cline from the Caucasus region to Russia Far East (Raghavan et al., 2015;de Barros Damgaard et al., 2018;Sikora et al., 2019). Ancient DNA study of 101 Bronze Age Eurasian ancient humans has found that large-scale population migration, admixture and replacement have reshaped the modern Eurasian demographic structure, including the westward spread of Yamnaya/Afanasievo populations into Central Asia, Xinjiang Tianshan mountain region in Northwest China and the Altai-Sayan region (Allentoft et al., 2015;Ning et al., 2019). Eastward expansion of the steppe pastoralists and their descendants gradually admixed or replaced by local indigenous steppe nomads and formed the multi-originated Scythian pastoralist tribes (Damgaard et al., 2018), as well as the later formed Xiongnu/Xianbei/Rouran/Uyghur/Turkic/Mongolic confederations. These eastern Eurasian nomadic pastoralist empires became the dominant groups and their subsequent westward migrations reshaped the genetic landscape of populations from the Eurasian steppe once again (Damgaard et al., 2018;Jeong et al., 2020). Modern Eurasian population genomic history also documented the large-scale western-eastern Eurasian admixture and massive population movement (Yunusbayev et al., 2015). Focused on language diversity, modern populations belonging to Indo-European, Uralic, Tungusic, Mongolic, and Turkic families/groups have widely distributed in Eurasia (Wang and Robbeets, 2020). From the cultural perspectives, significantly different steppe-pastoralist-associated cultures subsequently replaced or mixed with the existing incoming contemporaneous cultures, such as the cultural evolutionary sequence of the Yamnaya, Afanasievo, Sintashta, Andronovo, and other historic cultures (Allentoft et al., 2015;Damgaard et al., 2018;de Barros Damgaard et al., 2018). However, the extent to which these genetically/historically/archeologically attested shifts in culture, genetics, and language have shaped the genetic landscape of modern Chinese populations remains to be explored.

Recent ancient/modern genomes from China have provided some important new insights into the genetic formation of modern East Asians (Wang et al., 2020b). Yang et al. reported twenty-six Neolithic-to-historic genomes from Shandong, Fujian and the surrounding regions and found that the modern-DNA-attested North-to-South genetic differentiation has existed in East Asia since the early Neolithic period. North-South bidirectional migration and coastal continental East Asian population movement have shaped the observed East Asian genetic variation (Yang et al., 2020). Wang et al. reported one large-scale ancient DNA study focused on ancient human remains from South Siberia and Mongolia in the north to Taiwan in the south, which documented a strong culturally/genetically western-eastern communication in the contact region of the eastern Mongolian steppe and reported three large Holocene population migrations respectively from Northeast China, Yellow River basin and Yangtze River basin, and these expansion events were partially or completely associated with the spread of the main language families existing in East Asia (Wang et al., 2020a). Ning et al. sequenced genomes from the northern Chinese Neolithic-to-Iron-Age populations and documented the strong association between the subsistence strategy changes and population transformation (Ning et al., 2020). Ancient genomes of Shirenzigou Iron Age people have documented the western Yamnaya ancestry in Northwest China (Ning et al., 2019). However, the potential influence of western Eurasian ancestry in modern northwestern Chinese populations in molecular anthropology, medical genetics, and precision medicine kept unknown.

The development and prosperity of commercial and trade exchanges in the historically documented Silk Road further facilitated the recent eastern-to-western communication, however, the extent to which this cultural exchange or prehistoric cultural transformation in Northwest China was accompanied by population migration or only adoption of idea needed to be formally tested. The formation of the Hui people and their Muslim culture spread was one of the most important cases for exploring and testing these two opposing hypotheses. Demic diffusion model stated that all modern East Asian Muslim culture originated in western Eurasia and spread into China with a substantial external gene flow. However, the cultural diffusion model stated that the eastward dissemination of Muslim culture did not accompany by the large-scale population movement. Scholars have conducted several population genetic surveys based on the STRs, InDels, or other lower-density genetic markers, and they found that modern Chinese Hui originated from East Asia. However, others hold the opposite opinion that East Asian Hui people descended from western Eurasian migrants (Wang et al., 2019). Zhou et al. genotyped 30 InDel loci in 129 Ningxia Hui and found that East Asian populations contributed more genetic variants into Hui people compared with western Eurasian populations (Zhou et al., 2020), which was consistent with Zou’s observation in the Wuzhong Hui population (Zou et al., 2020). He at al. sequenced 165 ancestry informative Single Nucleotide Polymorphisms (AISNPs) in 159 Hui people and found East Asian-dominant ancestry in Hui along the Silk Road, which was in accordance with the autosomal STR-based results (Yao et al., 2016;He et al., 2018). However, the Y-SNP-based population genetic survey also identified western Eurasian founding lineages in East Asian Hui (Wang et al., 2019). Thus, the whole-genome-based genetic investigation was needed to be conducted to explore the origin, diversification, and subsequent genetic admixture of modern East Asian Hui people.

Sichuan is located in the interior of southwestern China is one of China’s 23 provinces and its capital which is bounded by 26°03′∼34°19′ north latitude and 97°21′∼108°12′ east longitude with Shaanxi, Gansu and Qinghai in the north, Tibet in the west, Yunnan and Guizhou in the south and Chongqing in the east. Its census population size over 91.29 million, and all Chinese officially recognized ethnic populations have been lived here. Hui population with a population size over 10.86 million. Sichuan Hui mainly distributed in many areas of the northwestern, southern, northern and eastern Sichuan with a population size over 0.11 million. Here, we obtained high-density SNPs data in 49 Hui people and 60 neighboring Han individuals and made one of the most comprehensive population comparisons based on the co-analysis of genome-wide data among all available modern and ancient Eurasian reference populations (Jeong et al., 2020;Ning et al., 2020;Wang et al., 2020a;Yang et al., 2020). We aimed to solve the following two questions: I) determine the extent of genetic heterogeneity or homogeneity among geographically different Hui people or between Hui and their adjacent neighbors; II) explore the timing, admixture sources and origin of modern Hui and to validate which models (Cultural diffusion versus demic diffusion) played the key role in the formation process of modern Sichuan Hui.

## Material and Methods

### Sample collection and DNA preparation

We collected saliva samples in 60 Nanchong Han and 49 Boshu Hui individuals in Sichuan province, Southwest China (**Figure S1**). Each included individual was indigenous people living there at least three-generation and the offspring of non-consanguineous marriage within Hui or Han. We collected all samples with the written informed consent and giving the corresponding genetic feedback of genetic testing focused on their ancestral composition and genetic health status. This project was inspected and approved by the Medical Ethics Committee of the North Sichuan Medical College. Our studied protocol was also followed the recommendations stated in the Helsinki Declaration of 2000 (Association, 2001). Genomic DNA was isolated via the PureLink Genomic DNA Mini Kit (Thermo Fisher Scientific) and preserved at −4°C until the next amplification.

### Genotyping, quality control, data preparation and two working datasets

We used Infinium® Global Screening Array (GSA) to genotype approximately 6,992,479 SNPs from autosome (645,199), Y-chromosome (26,341), X-chromosome (22,512) and mitochondrial genomes (4198). Genotype calling was carried out followed the default parameters. PLINK 1.9 (Chang et al., 2015) was used to filter out the raw genotype data and only remained markers with the site missing rate per person or missing rate per SNP below 0.01 (mind: 0.01 and geno: 0.01). Generally, the final dataset kept the autosomal SNPs and samples consistent with the following criteria: I) initial dataset only remained samples with the genotyping success rate larger than 0.99, II) p-value of the Hardy-Weinberg exact test larger than 0.001 (--hwe 0.001), III) SNPs with minor allele frequency of >0.01 (--maf 0.01) and a genotyping success rate per SNP higher than 0.01. We merged our newly-generated 109 genomes with previously published modern and ancient population data included in the Human-Origin dataset (lower-density dataset with more modern reference populations, including 72,531overlapping SNPs) and 1240K dataset (higher-density dataset with 193,838 overlapping SNPs) from Reich Lab (https://reich.hms.harvard.edu/downloadable-genotypes-present-day-and-ancient-dna-data-compiled-published-papers) or recent publications (Ning et al., 2020;Wang et al., 2020a;Yang et al., 2020). Genome-wide data of geographically different Hui from Guizhou also included in our formal admixture analysis(Wang et al., 2020c).

### Principal component analysis

We carried out the principal component analysis (PCA) using the smartpca package built-in EIGENSOFT (Patterson et al., 2012) with the default parameters based on the LD pruned and MAF filtered dataset. We also added two additional constraints (numoutlieriter: 0 and lsqproject: YES) to project ancient populations onto the PCA background. We performed two sets of PCAs based on different datasets: one East Asian-based PCA focused on 1325 individuals among 158 populations, and one Southern East Asian-based PCA focused on 963 individuals among 107 populations.

### Pairwise Fst genetic distances

We calculated Fst genetic matrix (Weir and Cockerham, 1984) using our in-house script and PLINK 1.9 (Chang et al., 2015) to explore the genetic similarities and differences between Boshu Hui, Nanchong Han and other Eurasian modern and ancient populations.

### ADMIXTURE clustering

Using PLINK 1.9 (Chang et al., 2015), we removed the strongly linked SNPs with the R-value larger than 0.4. We used 200 SNPs as the windows width and 25 SNPs as the sliding window with the following parameters (--indep-pairwise 200 25 0.4). We run the model-based genetic clustering using ADMIXTURE 1.3.0 (Alexander et al., 2009) with the unsupervised mode. We used the predefined ancestral populations ranging from 2 to 20 (K: 2∼20) with 10 replicates and found K=13 with the best resolution for ancestry dissection (**Figure S2**).

### Allele-based shared ancestry estimation

We used the *qp3pop* packages in ADMIXTOOLS (Patterson et al., 2012) to perform the *outgroup-f*_*3*_*(Source1, Source2; Mbuti)* to evaluate the shared genetic drift among 366 Eurasian modern and ancient populations using the default parameters. Followingly, we used the *qp3pop* package (Patterson et al., 2012) to perform the *admixture-f*_*3*_*(Source1, Source2; Boshu Hui/Nanchong Han)* to explore the admixture signatures with different Eurasian ancestral source candidates, where significant negative-*f*_*3*_ values with Z-score less than −3 denoted that the targeted population was the admixture result between two parental populations of source1 and source2. We finally used the *qpDstat* package (Patterson et al., 2012) to estimate the *f*_*4*_*-statistic* value with one additional parameter (*f*_*4*_Mode: YES), and standard errors were estimated using the block jackknife (Patterson et al., 2012).

### Phylogenetic relationship reconstructions

We used Neighbor-joining (NJ) algorithm to reconstruct the phylogenetic relationships based on different genetic distance matrixes in Mega 7.0 (Kumar et al., 2016) or TreeMix (Pickrell and Pritchard, 2012). We first built one NJ tree based on the invert *f*_*3*_-based genetic distances (1/*outgroup-f*_*3*_) among 366 Eurasian populations to explore the genetic relationship between modern and ancient populations and newly focused populations and other reference groups. We following run TreeMix (Pickrell and Pritchard, 2012) with the default settings and 100 replicates to choose the best-fitted model and explore the phylogenetic relationship among 47 modern populations with the migration events ranging from 0 to 11.

### Modeling admixture history

We run *qpWave/qpAdm* packages in ADMIXTOOLS (Patterson et al., 2012) to explore the minimum ancestral population number via the rank tests and then evaluated the corresponding admixture ancestral proportion. We used one additional parameter of ‘allsnps: YES’, and nine right reference outgroup populations were used here (Mbuti, Ust_Ishim, Kostenki14, Papuan, Australian, Mixe, MA1, Onge, Atayal).

### Deep population history reconstruction

We used *qpGraph* (Patterson et al., 2012) to successfully-fit two admixture graph models to explore the observed genetic variations in Han and Hui populations. The following parameters were used: blgsize: 0.05; lsqmode: NO; diag: 0.0001; hires: YES; initmix: 1000; precision: 0.0001; zthresh: 0; terse: NO; useallsnps: NO. We followed two basic models to reconstruct the population genomic history of our targeted populations (Green et al., 2010;Reich et al., 2010;Wang et al., 2020b). The first skeletal framework was used from our previously reported model (Wang et al., 2020b), but we replaced the proxy of the northern Yellow River farmer as a more geographically close millet group. The second model was reconstructed based on the two archaic introgression events among non-Africans (Green et al., 2010;Reich et al., 2010).

### Method for dating admixture events

We used Admixture-induced Linkage Disequilibrium for Evolutionary Relationships (ALDER) (Loh et al., 2013) to estimate the date-scale of population admixture events for two Hui populations (Boshu and Guizhou) and Nanchong Han based on two datasets and the default parameters. First, we employed 28 potential ancestral candidates (Twenty-five eastern Eurasian-like source: Htin_Mal, Kinh_Vietnam, Mlabri, Ami, Atayal, Dao, Hmong, PaThen, Mongol_Uuld, Daur_HGDP, Han_Shandong, CoLao, Dai_HGDP, Gelao_Longlin, LaChi, Li_Hainan, Maonan_Huanjiang, Zhuang_Guangxi, Tibetan_Lhasa, Tibetan_Shigatse, Xijia_Kaili, Dongjia_Kaili, Gejia_Kaili, Ulchi, Nganasan; Three western Eurasian-like source: French, Greek, Basque) included in the merged Human Origin dataset to date the admixture times for Han and Hui. We also used the Nanchong Han as the East Asians and other western Eurasians to date the formation of the Boshu Hui people. Secondly, we used we also fourteen populations (Dai, Altaian, French, Mongolian, Oroqen, Tajik, Ami, Sardinian, Greek, Atayal, English, Pathan, Basque, Han_Nanchong) included the merged 1240K dataset to date the admixture times of Chinese Hui. We calibrated the years used the following formula: Year=1950-28*(Generation-1). To validated the concordance of our results, we also estimated the admixture times of Uyghur populations, our observed admixture patterns were consistent with previous reports (Loh et al., 2013).

### Y-chromosomal and mtDNA haplogroup assignment

We followed the recommendations of ISOGG and used our in-house script to assign the Y-chromosomal paternal lineage and used the HaploGrep 2: to assign mitochondrial maternal haplogroups (Weissensteiner et al., 2016).

## Results

### General patterns of genetic structure

We generated the whole-genome SNP data from 109 western Chinese Hui or Han individuals in Sichuan province and co-analyzed with the publicly available modern and ancient DNA data in the comprehensive population genetic history reconstruction. We grouped modern reference individuals from Eurasia according to their language family categories (including Altai or Tans-Eurasian (Turkic, Mongolic, Tungusic, Japonic, and Koreanic), Hmong-Mien, Tai-Kadai, Austronesian, Austroasiatic, and Sino-Tibetan (Sinitic and Tibeto-Burman)) and classified ancient peoples based on their geographical affinity (**Figure 1**). We first carried out the PCA to explore the broad overview of the population structure of all modern eastern Eurasians and ancient individuals were projected on the top two PCs axes (**Figure 1A**). This inference revealed four genetic clusters (northeastern Tungusic/Mongolic cluster, southwestern Tibeto-Burman cluster, southern inland Hmong-Mien cluster and southern Austronesian/Austroasiatic cluster) and one Sinitic-Tai-Kadai cline. Boshu Hui was clustered with northern Han Chinese and other Northern East Asian minorities, and Nanchong Han were clustered dispersedly and formed the clines partially overlapped with previously published Hans. We also found a close genetic affinity between Boshu Hui and late-Bronze-Age-to-Iron-Age Northern East Asians from Henan province (Haojiatai and Luoheguxiang). In the Southern East Asian-based PCA, we observed clear population substructure, such as Hmong-Mien and Sino-Tibetan genetic lines separated from others (**Figure S3**). We expected to observe a unique genetic cluster or cline of Hui if this studied Hui possessed unique western Eurasian-originated ancestry, however, we found that studied Han and Hui were localized in the intermediate position between the highland East Asians and Southern East Asians.

**Figure 1.**
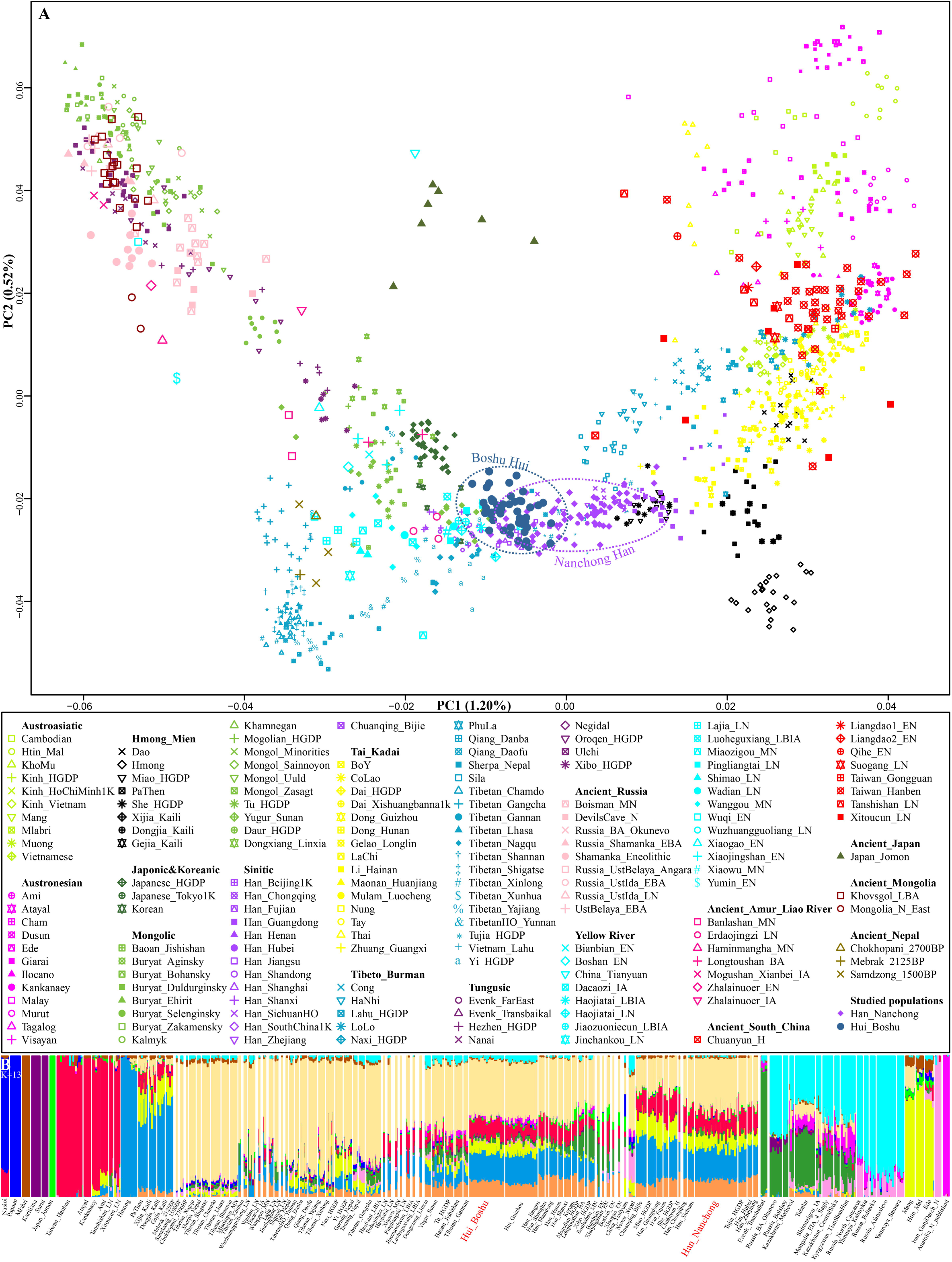
Overview of population structure. **(A)** Principal component analysis (PCA) based on the top two components of East Asian genetic variations. Included ancient individuals were projected onto the modern genetic background. **(B)** Ancestry component composition between Boshu Hui, Nanchong Han and other Eurasian reference populations when the optimal K value equal to thirteen.

Model-based ADMIXTURE results with the optimal K value (k=13) have identified multiple East Asian dominant ancestry which were enriched in the both inland (Hmong) and coastal (Ami) Southern East Asians, and others were maximized in the inland (Tibetan) and coastal (Japanese) Northern East Asians (**Figures S2 and S4**). Our studied present-day western Hui shared most of their genetic ancestry related to Northern East Asians (Tokyo-Japanese-like: 0.123 and Mebrak_2025BP-like: 0.398) and Southern East Asians (Hmong-like: 0.208, Taiwan_Hanben-like: 0.116 and Htin_Mal-like: 0.082, (**Figure 1B** and **Table S1**). We also found ancestry related to western Eurasian (Russia_Poltavka: 0.008). Adjoining Nanchong Han possessed dominant Northern East Asian ancestry (Japanese: 0.130, and Mebrak: 0.344) and Southern East Asian ancestry (Hmong: 0.230, Htin: 0.116, and Hanben: 0.140). The pattern of genetic relationship revealed by model-based ADMIXTURE was consistent with the PCA results.

### Stronger genomic affinity with Northern East Asians

To quantitatively evaluate the genetic differences between Hui and Han from Sichuan province and other 170 modern and 67 ancient Eurasian populations with a population size larger than five, we calculated the pairwise Fst genetic distance (**Table S2**) according to Weir and Cockerham (Weir and Cockerham, 1984). Fst-based comparison of genetic differences showed that Chinese Hui was most closely with geographically close northern Han populations, such as Han from Nanchong (0.0019) and Beijing (0.0013). Among ancient populations, Boshu Hui had the close genetic relationship with Mongolia_EIA_1_SlabGrave (0.0603), followed by Russia Shamanka Eneolithic (0.0675), Mongolia_LBA_4_CenterWest (0.0680), and it had a relatively larger genetic distance with Yellow River Yangshao farmer (Wanggou_MN: 0.1557) and Yangtze rice farmer (Xitoucun_LN: 0.1729). A subtle different pattern of genetic affiliation of Nanchong Han and their eastern Eurasian modern and ancient reference populations was identified in the matrixes of Fst. Nanchong Han tended to shift a close relationship with central and southern Han Chinese (Zhejiang Han: 0.0003, Hubei Han: 0.0005 and Sichuan Han: 0.0006).

We subsequently used a formal test of outgroup-*f*_*3*_-statistics in the form *f*_*3*_*(Test populations, Hui_Boshu/Han_Nanchong; Mbuti)* to explore the shared genetic drift between two studied populations and Eurasian reference populations, included 188 modern and 176 ancient populations, which may be contributed to the genetic materials into the gene pool of Hui and Han in Sichuan province (**Table S3** and **Figures 2, 3A∼B and S5**). We found that Boshu Hui possessed the most shared ancestry with modern Han groups from Hubei (0.3004), Fujian (0.2992) and Henan (0.2992) provinces and ancient Iron Age Luoheguxiang (0.2989) and late Neolithic Pingliangtai (0.2978) after excluding some groups with the bias introduced via sample size or batch effects. Nanchong Han shared the most genetic drifts with modern Han groups from Hubei (0.3031), Fujian (0.3025) and Chongqing (0.3020) provinces and ancient Iron Age Luoheguxiang (0.3013) and late Neolithic Pingliangtai (0.3004). Small shared genetic drifts between archaic people and studied groups (Denisovan: 0.0217 in Han and 0.0224 in Hui; Neandertal: 0.0249 in Han and 0.0251 in Hui) were identified. There is a strong statistically correlation of shared ancestry between Hui and Han in the outgroup-*f*_*3*_ values (*f*_*3*__Han = 0.9579\**f*_*3*__Hui + 0.0099; R^2^ = 0.9992). Correlation efficient between shared genetic drift with the differences in latitude (0.1817 in Han and 0.1691 in Hui) and longitude (0.6992 in Han and 0.6777 in Hui) showed the association between the shared genetic drift and the longitude, which was consistent with the gradient changes in the *f*_*3*_-based heatmap (**Table S3**).

**Figure 2.**
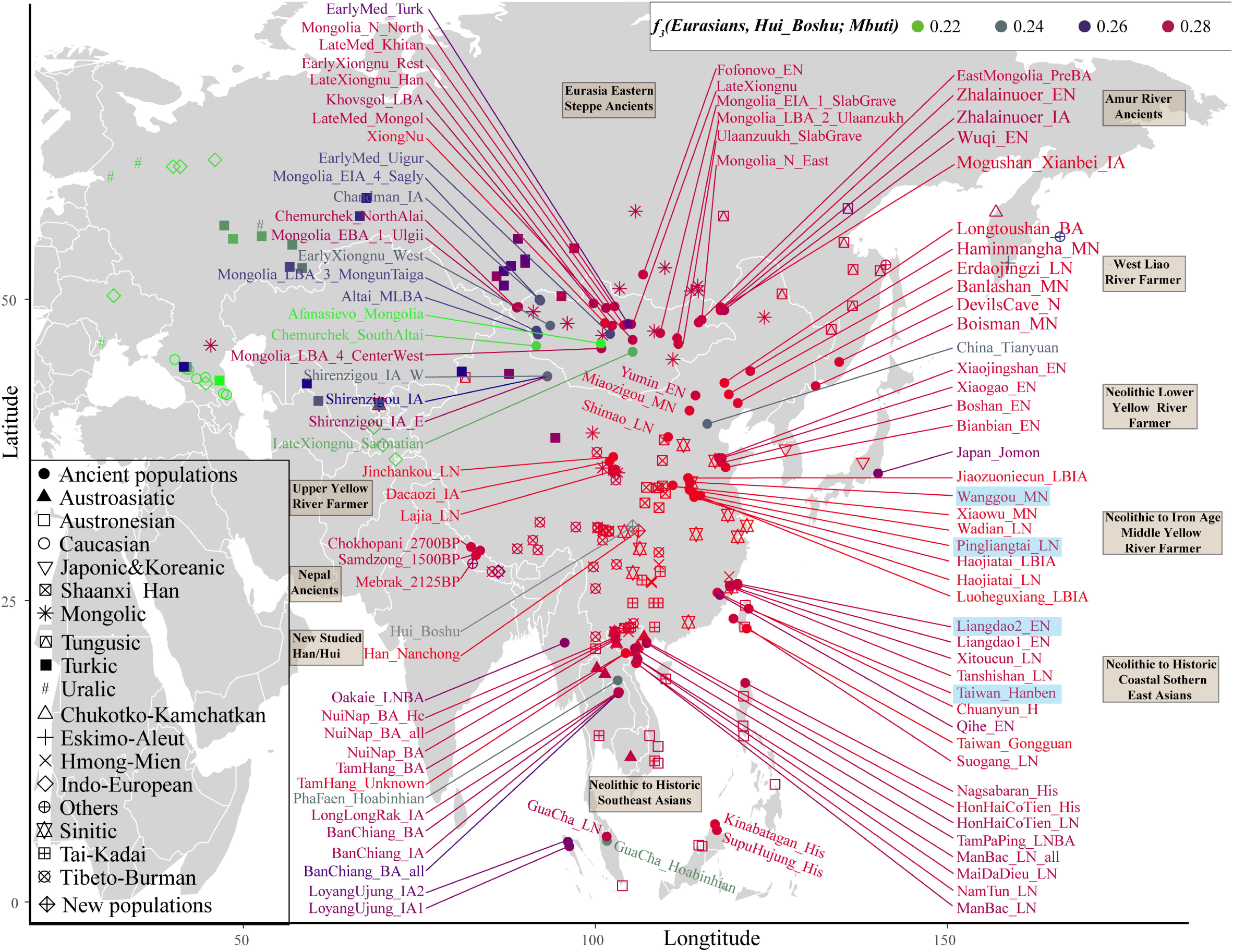
Heatmap showed the genetic affinity between Boshu Hui and modern/ancient Eurasians estimated by *f*_*3*_-statistic in the form *f*_*3*_*(Eurasian, Hui_Boshu, Mbuti)*. Red color donated higher genetic affinity, suggested their genetic to Hui people. The green color denoted a smaller *f*_*3*_-value, suggested their distant relationship with the Hui people. Here, the main ancient populations were used in the qpGraph-based admixture modeling were marked with the light blue background. LBA: Late Bronze Age, IA: Iron Age, MLBA: Middle and Late Bronze Age, LN, Late Neolithic, EN, Early Neolithic, MN, Middle Neolithic, LBIA: Late Bronze Age and Iron Age.

**Figure 3.**
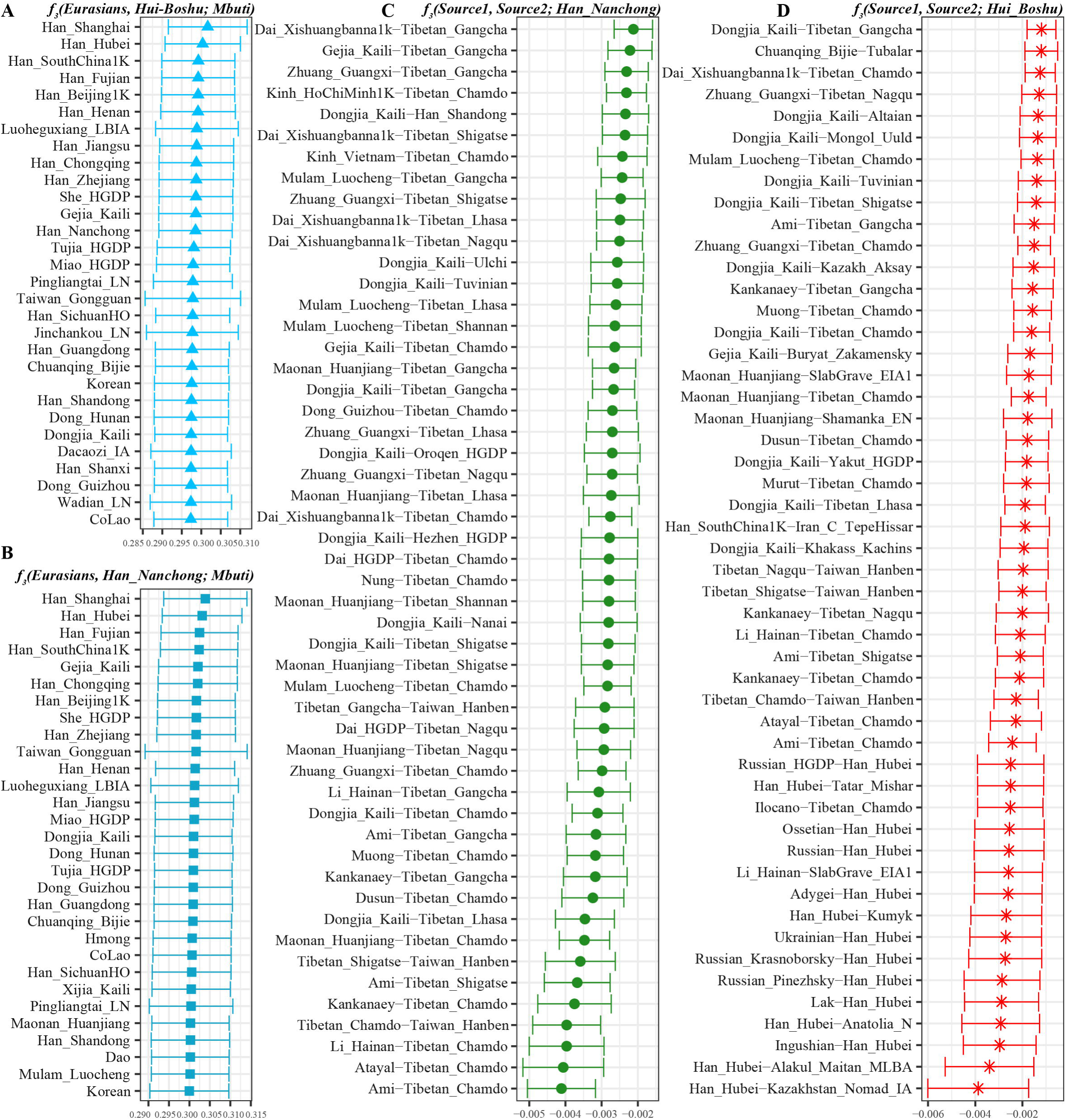
The shared genetic drift and admixture signal for the targeted Han and Hui populations. (**A∼B**). The top thirty populations harboring genetic affinity with Boshu Hui (**C**) and Nanchong Han (**D**). (**C∼D**). Top sixty pairs of source populations possessing admixture signals via three-population testing in the form of *f*_*3*_*(Source1, source2; Boshu Hui/Nanchong Han)*. All showed population pairs with Z-scores less than negative three.

We also explored the potential ancestral sources of Hui and Han among these reference datasets via admixture-*f*_*3*_-statistics in the form *f*_*3*_*(Source1, source2; Hui_Boshu/Han_Nanchong)*, which would be expected to be obtained statistically negative values (Z-scores less than −3) when the allele frequency of Hui/Han has localized the intermediate position between source1’s frequency and source2’s frequency. We found signals supporting the two-way or three-way admixture models for the genetic formation of Sichuan Hui and Han (**Tables S4∼5**). North-South admixture signatures were identified when we treated Nanchong Han as the targeted population (**Table S4** and **Figure 3C**). Here, we could also identify genetic contributions from central and southeastern Han Chinese into Sichuan Han via the negative values in *f*_*3*_*(Han_Fujian/Guangdong/Hunan, other sources; Han_Nanchong)*, suggesting the possible westward of Han Chinese in the historic time. Focused on Chinese Hui, we obtained the most admixture signals among the source pairs of Northern East Asians (Tibetan and southern Siberian) and southern indigenous East Asian (Tai-Kadai and Austronesian speakers), such as *f*_*3*_*(Ami, Tibetan_Chamdo; Hui_Boshu)* = − 7.216*SE (standard error). Western Eurasian admixture signals could be also obtained when we used the Han Chinese as one source and Ingushian (−5.748), Lak (−5.508) or Adygei (−5.429) as the other source (**Table S5** and **Figure 3D**).

Focused on genetic homogeneity or heterogeneity between Hui and their neighbors, we performed *f*_*4*_-statistics in the form *f*_*4*_*(Han_Nanchong, Hui_Boshu; Eurasians, Mbuti)*. Here, we observed relatively smaller Z-scores, especially for negative Z-scores, thus we set the statistically significant threshold of absolute Z-scores as two (Table 1). Significant differences in the degree of allele sharing with the modern and ancient reference East Asians were observed due to the observed significant negative (stronger affinity with Hui) or positive (stronger affinity with Han) values, suggesting their differentiated demographic traces (**Table 1 and Tables S6∼7**). We documented gene flow between modern/ancient western Eurasian and Chinese Hui though significantly more allele sharing of western pastoralists with Hui than with Nanchong Han (**Table 1**), and also documented strong East Asian affinity of Han compared with Hui. We further identified close genomic affinity between Hui and geographically close Han via significant negative *f*_*4*_-values in *f*_*4*_*(Eurasian, Han_Nanchong; Hui_Boshu, Mbuti)* based on two merged datasets (**Tables S8∼9**). Similarly, we performed these types of *f*_*4*_-statistics (**Table S10**) to explore the genetic homogeneity between Guizhou Hui and Boshu Hui. Guizhou Hui harbored excess sharing alleles related to western steppe pastoralists than Boshu Hui. The identified differentiated population structure showed their different demographic history, Thus, more density geographically distinct Hui populations should be collected and genotyped to comprehensively characterize the landscape of East Asian Hui.

Previous genetic studies have demonstrated that the historically/archaeologically attested spread of millet and rice agriculture in East Asia was accompanied by the bidirectional spread of farmers from the northern Yellow River basin and southern Yangtze River basin (Yang et al., 2020). As shown in **Figure 4A** and **Table S11**, all statistically positive *f*_*4*_-values observed among modern Eastern panels based on the merged 1240K dataset in *f*_*4*_*(Han/She/Tujia/Ami/Atayal/Miao/Naxi/Yi, Eastern Eurasian; Hui_Boshu, Mbuti)* and ancient Northern East Asian panels in *f*_*4*_*(Haojiatai_LN/Jiaozuoniecun_LBIA/Pingliangtai_LN/Luoheguxiang_LBIA/Xiaowu_MN/Erdaojingzi _LN/Wadian_LN/Dacaozi_IA/Shimao_LN/Jinchankou_LN/Wanggou_MN/Taiwan_Gongguan/Haojiatai_LBIA/Longtoushan_BA, Eastern Eurasian; Hui_Boshu, Mbuti)* demonstrated that Chinese Hui harbored a stronger East-Asian-like, especially Northern East Asian-like ancestry. To further test which populations directly contributed more ancestry into Hui populations, we first performed *f*_*4*_*(Eurasian reference, Hui_Boshu; potential ancestral source, Mbuti)*. Most significant negative *f*_*4*_-values were observed when we used modern populations (Ami, Atayal, Dai, Han, Thai, Kinh, Lahu, She, Miao) and ancient Taiwan Hanben people as the potential ancestral source candidates (**Figure S6** and **Table S12**). We further found that Neolithic-to-Iron-Age populations from the Yellow River basin shared more derived alleles with Boshu Hui relative to modern East Asians and ancient Southern East Asians, which pointed to the stronger Neolithic-to-modern Northern East Asian affinity of modern Hui. To further study other additional gene influx into Hui compared with Northern East Asian sources, we performed symmetrical-*f*_*4*_*(Ancestral sources, Hui_Boshu; other Eurasian reference populations, Mbuti)*. Here, significant negative *f*_*4*_-values indicated additional genetic contributions from the reference group relative to the northern ancestral source, and non-significant *f*_*4*_ results denoted that Hui was the direct descendant of northern sources (**Table S12**). These genetically attested patterns of genetic admixture signatures based on the 1240K dataset are further confirmed with the merged Human Origin dataset (**Figure 4B, Figure S7** and **Tables S13∼14**). Results from *f*_*4*_-statistics in the form *f*_*4*_*(Reference population1, reference population2; Hui_Boshu, Mbuti)* documented gene flow between modern Sinitic-speakers and Hui based on significantly more allele sharing of Hubei Han/other speakers with Hui than to other modern and ancient Eurasians. Interestingly, we found that the middle Yellow River farmers formed one clade with Boshu Hui when it compared with modern East Asians, but we could still identify negative *f*_*4*_-values when we used lower or upper Yellow River ancients and other northern ancients as the potential sources (Wadian_LN, Lajia_LN, Wuqi_EN, Xiaojingshan_EN, Shimao_LN, Bianbian_EN, Xiaogao_EN, Longtoushan_BA, Zhalainuoer_EN, Banlashan_MN and Boshan_EN) comparing to modern East Asian or Southern East Asian Hanben, pointing to additional gene flow from Southern East Asians (**Figure 4B**).

**Figure 4.**
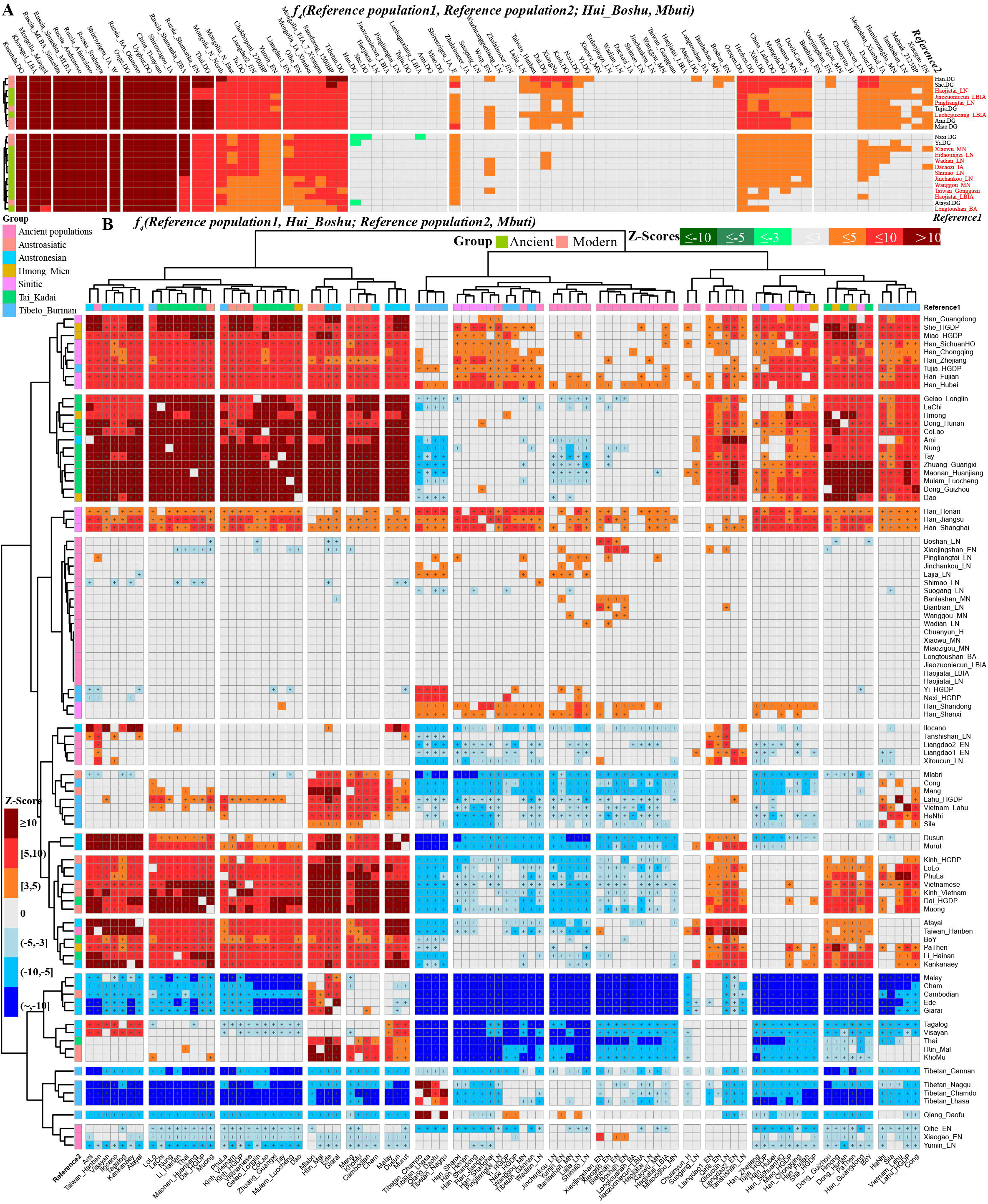
Excess of sharing alleles between Boshu Hui and modern and ancient Eurasians. **(A)** Symmetrical *f*_*4*_-statistics in the form *f*_*4*_*(Reference population1, reference population2; Hui_Boshu, Mbuti)* to test their genetic affinity between Eurasian reference populations based on the merged 1240K dataset. Red color showed Boshu Hui shared more alleles with Northern East Asians (Right population lists). **(B)** Affinity-*f*_*4*_-statistics in the form *f*_*4*_*(Reference population1, Hui_Boshu; reference population2, Mbuti)* to explore the genetic continuity and admixture between potential ancestral sources and Boshu Hui based on the merged Human Origin dataset. The red color denoted the third population (Bottom population lists) shared more derived alleles with the first population (Right population lists) compared with the second one. And the green color denoted the third population shared more alleles with the second one relative to the first one. Statistically significant f-statistics were marked as “+”.

**Figure S8∼10** and **Table S11∼14** also focused on the genomic history of geographically adjacent Han Chinese, we found that more modern and ancient East Asian populations shared more alleles with Nanchong Han than other reference groups do, as shown by the positive *f*_*4*_ values presented in **Figures S8A and S9**. Symmetrical-*f*_*4*_-statistics focused on the Nanchong Han in **Figure S8B and S10** further demonstrated that Nanchong Han shared more Northern East Asian-like derived alleles and simultaneously obtained the additional gene flow from Southern East Asians. Positive values of *f*_*4*_*(Han_Nanchong, Hui_Boshu; modern/ancient Southern East Asians, Mbuti)* also documented more Southern East Asian-like ancestry in Nanchong Han compared with Boshu Hui (**Table 1**).

### Genomic formation and population transformations of East Asian Hui and their neighbors

Considering our observed admixture events and sources of our studied Han and Hui, we applied *qpWave/qpAdm* to model their ancestry. *QpWave/qpAdm* was the formal test that could evaluate whether Han/Hui was consistent with the pattern of descending from one or more ancestral source populations relative to some chosen right or reference outgroup populations (being differentiated genetic relationship with the included source populations) and then evaluated their corresponding ancestral proportions. Results from *qpWave* showed the best two-way admixture model could be used to model the genetic formation of Han (p_rank1>0.05) and three-way admixture model for Hui (p_rank2>0.05). Modeling Nanchong Han as a mixture of ancestral Northern East Asians (Xiaowu, Pingliangtai, Luoheguxiang, Shimao and Miaozigou) and Southern East Asian (Hanben), we estimated that 0.697 (Shimao_LN-like) ∼ 0.879 (Pingliangtai_LN-like) of their ancestry is Northern East Asian-like, mostly associated with Yangshao and Longshan people from Henan province (**Table S15** and **Figure 5A**). Higher Northern East Asian ancestry proportion could be modeled when we used the populations in the later periods (late Neolithic to historic) and geographically close populations from Central Plain in North China. The Southern East Asian Hanben-like ancestry proportion spanned from 0.121 to 0.303 in different two-way admixture models. These patterns of admixture processes were consistent with the North China Origin hypothesis of modern Sinitic speakers and continuously absorbed gene flow from the Southern East Asians. We found that most of the ancestry in Hui’s gene pool was derived from East Asians, especially from Northern East Asian. The best-fitted three-way admixture proximal models for Boshu Hui as the mixtures of major ancestry from northern and Southern East Asians and additional 0.019∼0.051 of its ancestry is ancient western Eurasian Andronovo-like or 0.021∼0.052 of its ancestry associated with modern French. We finally modeled Hui’s gene pool via a two-way admixture model used geographically close Nanchong Han as an East Asian source and five modern European and twelve western Eurasian steppe pastoralist populations as the western Eurasian source (**Table S16**). Hui can be modeled as the admixture product of 0.023∼0.033 ancestry related to western Eurasians and 0.967∼0.977 ancestry related to Nanchong Han with different source pairs.

**Figure 5.**
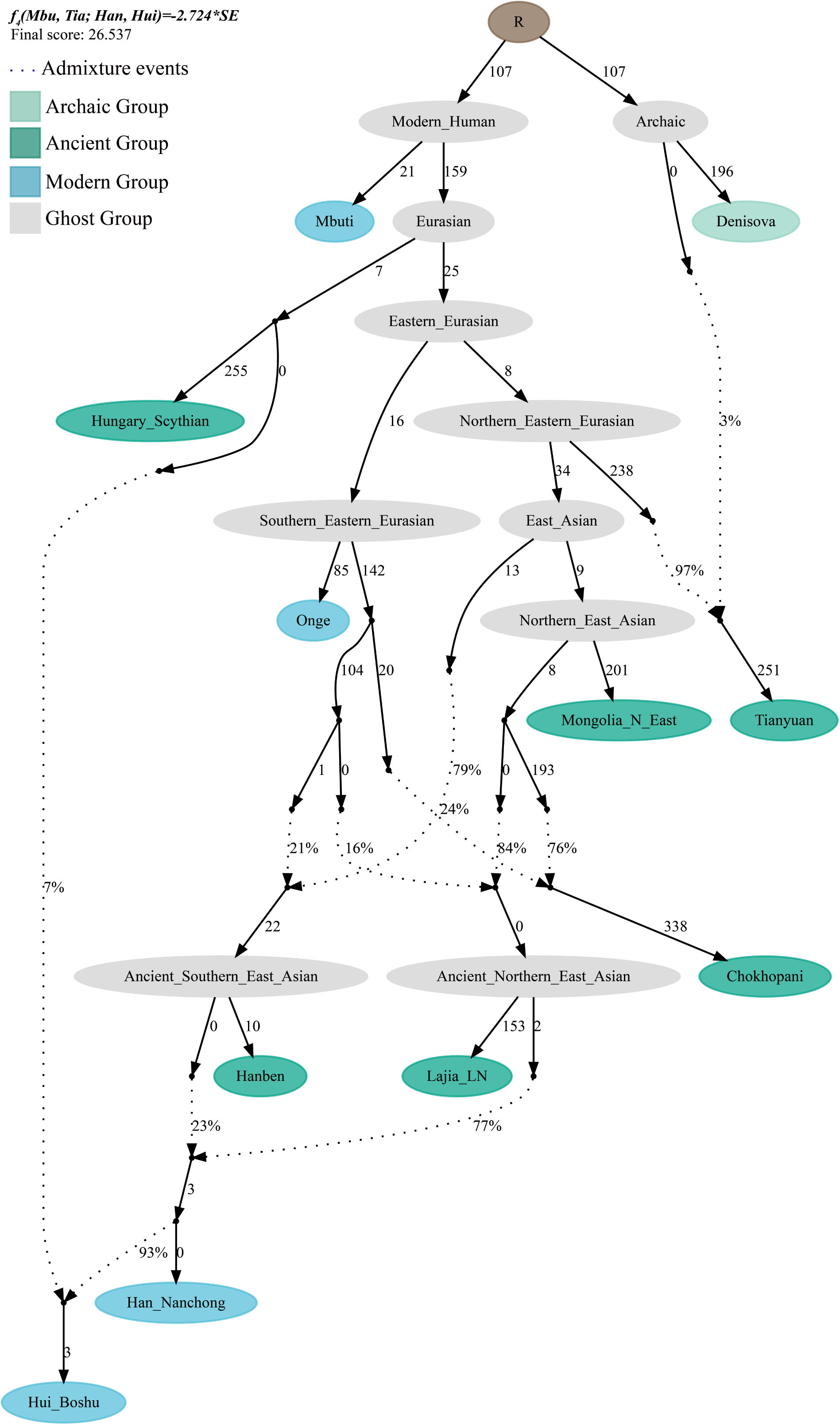
*qpAdm*-based ancestry composition and *qpGraph*-based admixture graph. **(A)** *QpAdm* result showed the two-way admixture for Nanchong Han and the three-way admixture model for Boshu Hui. Red Bar showed standard error. **(B)** *QpGraph*-based phylogeny showed the western Eurasian gene flow event in Boshu Hui. Branch length was marked with the *f*_*2*_ shared drift distance (1000 times). Admixture events were denoted as dotted line. Admixture proportion was marked along the dotted line. Here, we used Hungary Scythian as the western source. **(C)** Complex *qpGraph*-based admixture graph displayed the two gene flow events into Nanchong Han. MNE: Mongolian N_East.

We next used *qpGraph* to fit the phylogeny-based model with orders of population splits, branch length measured by the *f*_*2*_ values, admixture events and corresponding ancestral proportions (**Figure 5**). We used modern South Asian indigenous Hunter-gather (One) and 40,000-year-old Tianyuan people as the early eastern Asian lineages, and used Mongolia Neolithic people (Mongolia_N_East), late Neolithic Qijia people (Qijia_LN) and Tibet Plateau ancient (Chokhopani) as the proxies of the northern East Asian lineages and Iron Age Hanben as southern East Asian proximity. We also used temporally different western Eurasians spanning from early Bronze Age to nowadays as the western sources. For western Han Chinese, we modeled it as the admixture of the major ancient northern East Asian ancestry related to the Lajia_LN (71∼80%) and the remaining ancestry from ancient southern East Asian related to the Hanben (20∼29%). We further successfully fitted a model with minor gene influx from western Eurasian into East Asian Hui people (approximately 7% related to Hungary Scythian). We further validated this model and confirmed the minor genetic contribution from western Eurasians with different temporally different central Asian or European sources (7% in Early Neolithic Bronze Age Kalmykia Yamnaya (**Figure S11**), 7% in Middle and Late Bronze Age Sintashta (**Figure S12**), 6∼8% in other Iron Age to modern western Eurasians (**Figures S13∼17**).

We further used the ALDER-based method to estimate the admixture date of the introduced western Eurasian ancestry into the gene pool of East Asian populations and found the genetic contact between western and eastern Eurasian. Firstly, we validated whether our merge datasets were suitable for dating admixture times via examining the admixture history of Uyghur which was previously genetically attested via admixture between French and Han in twenty generations ago (Loh et al., 2013). Our results also showed evidence of admixture for Uyghur (**Table S18**), about 17.9±1.65 generations (501.2 years) ago in the French-Nanchong Han model and 18.86±1.79 generations (528.08 years) ago in the French-Shandong Han model, which suggested that our merged datasets were suitable for estimating admixture process of Hui. Thus, we then model the admixture process of Hui used geographically close Han as the East Asian source and we found the admixture generations ranged from 18.08±2.62 in French-Han model to 25.35±7.27 in Tajik-Han model. For Guizhou Hui, we observed more recent admixture times ranging from 17.29±3.18 (Basque-Han_Nanchong) to 18.81±1.9 (French-Han_Nanchong) generations based on the merged 1240K dataset and spanned from 18.87±1.73 (Han_Nanchong-Greek) to 19.51±1.68 (Han_Nanchong-Basque) generations based on the merged Human Origin dataset. We further estimated the admixture times used different East Asian sources based on two datasets and we found that admixture events mainly occurred during the historic times ranging from 1314.12±103.6 CE (21.71±3.7 generations) to 1091.24±180.88 CE (29.67±6.46 generations) with Basque as the proxy of western sources or ranging from 1367.6±100.52 CE (19.8±3.59 generations) to 1120.64±238 CE (28.62±8.5 generations) with French as the proxy of western sources. Other detained admixture times for Guizhou Hui were presented in **Table S18** which also evidenced their more recent admixture process than Boshu Hui, such as 20.04±1.18 generations for Guizhou Hui but 22.09±4.57 generations for Boshu Hui in the Ami-French admixture model. We also observed admixture events with different northern and southern East Asian sources at different times (**Figure S14 and Table S18**). We should pay attention to this estimated time as the general time, which is late than the first genetic admixture time due to the continuous gene flow and admixture. We also observed the complex genetic admixture process between northern and southern East Asian in both Han and Hui.

To comprehensively inspect the overall genetic relationship and cluster patterns of Eurasians, we constructed a most-representative NJ tree among 190 modern and 176 ancient Eurasian populations and based on the invert *f*_*3*_-based genetic matrixes (*1/f*_*3*_). We found that the Neandertal and Denisovan were fitted at the root in the reconstructed phylogeny and also identified a genetic differentiation between eastern modern and ancient Eurasian populations (**Figure 6**). Ancient populations were clustered close with their geographically close modern populations, suggesting the long-term continuity of the primary ancestry in the genetic transformation: Southeastern Neolithic-to-Historic people clustered with modern Austronesian; late-Neolithic-to-Historic Southeast Asians grouped with modern Austroasiatic and Tai-Kadai; ancient Nepal Tibetans clustered with modern highland Tibeto-Burman; Northern East Asians from Yellow River basin grouped with modern Sinitic people, Siberian ancients mainly clustered with modern Mongolic, Tungusic and Uralic people, and western ancient Eurasians grouped with modern Indo-European or Turkic people. We found a close genetic relationship between Boshu Hui and Nanchong Han, and they followingly clustered with geographically close Sichuan Han. The close phylogenetic relationships between the studied populations and Eurasian modern reference populations were further confirmed via TreeMix-based phylogeny (**Figure 7**). Here, we could identify western gene flow events into the Kazakh population but not into Huis, which supported the strong genetic affinity between Boshu Hui and modern East Asians and the small amount of western Eurasian-like gene influx.

**Figure 6.**
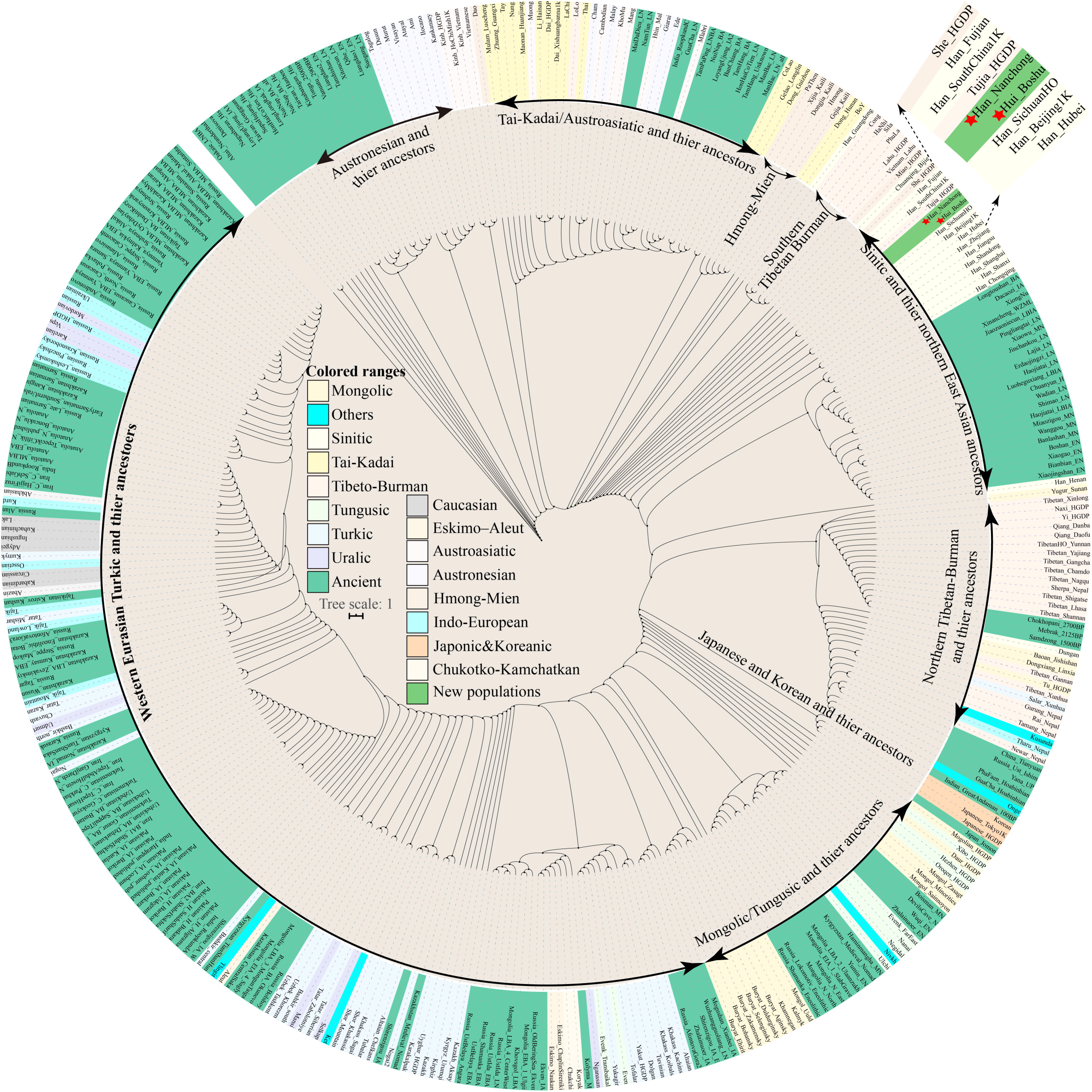
Phylogenetic relationship among Eurasian populations. The neighbor-joining tree was constructed based on the invert *f*_*3*_-based matrix (1/*f*_*3*_). All included populations were coded by different color backgrounds.

**Figure 7.**
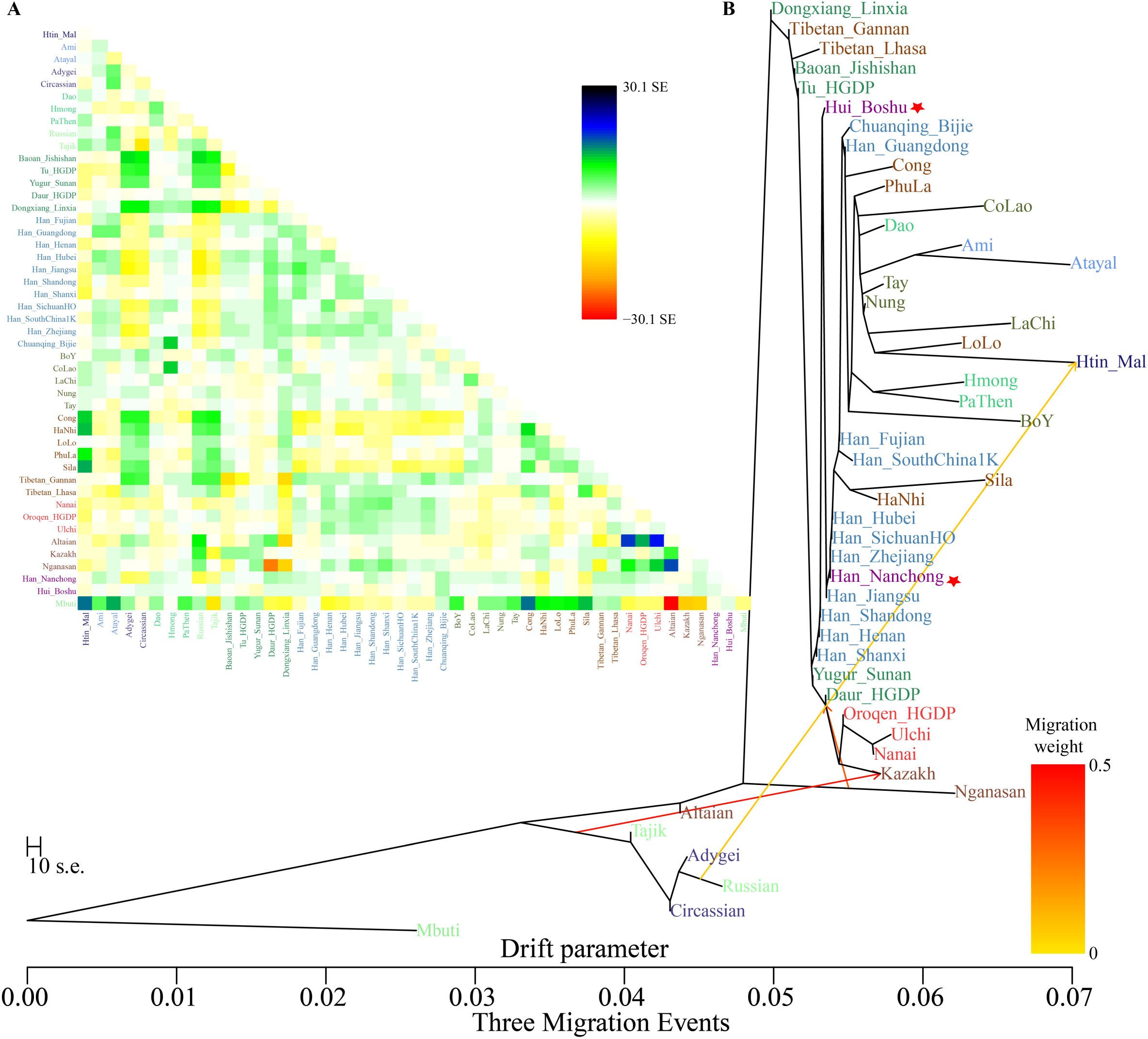
TreeMix results among 47 populations with three gene flow events. (A). Left upper heatmap showed the residual matrixes and (B) the right tree showed the population splits and admixture events. Populations were colored according to their language family categories.

### Paternal and maternal founding lineages in Han and Hui

We assigned 109 tested mitochondrial genomes based on 4,198 maternal mitochondrial lineage informative SNPs (MLISNPs) and 60 Y-chromosomal genomes (28 Huis and 32 Hans) based on 22,512 paternal Y-chromosomal lineage informative SNPs (YLISNPs). Among 49 studied Hui people (**Table S19**), we identified 36 maternal lineages with the terminal lineage frequency ranging from 0.0204 to 0.0816 (D4b2b, 4; R9b1b: 4). Maternal lineages of B4a, B4i1, D4a, F2, M7b1a1, M8a3a1 and F2d were also identified at least twice among the Hui population. 28 Hui males were assigned to 13 different terminal lineages with a frequency ranging from 0.0357 to 0.5357 (O2a2b1a2a1a3, 15). Here, we also identified some samples with Siberian-dominant paternal lineages (C1b1a2b, C2c1a1, C2c1b2b2∼ and R1b1a1a2a2c1a). For the studied Hans, we identified 51 different maternal lineages with the frequency ranging from 0.0167 to 0.0500 (M7b1a1e1, 3). We obtained 25 terminal paternal lineages among 32 males with the frequency ranging from 0.0323 to 0.0968 (N1b2a2∼, 3). We could also identify Siberian-derived lineages in Han populations (C2c1a1a1a, C2c1a2a2, C2c1a2b2, C2c1b2b2∼ and Q1a1a1a1a∼)

## Discussion

Two well-known hypotheses have been proposed to explain the diffusion processes of some special cultures or population movements: the cultural diffusion model and the demic diffusion model (Wen et al., 2004). Many cases of the demic diffusion model have been reported via the ancient and modern DNA studies, such as Bantu expansion in Africa (Patin et al., 2017), barley/wheat agriculturalists spread into Europe from the Fertile Crescent in Near East, and steppe nomadic pastoralist widely disseminated in the Eurasian steppe (Lazaridis et al., 2014;Narasimhan et al., 2019). In East Asia, ancient DNA studies have documented that the spread of the rice/millet agriculture, as well as corresponding language (agriculture-language-co-dispersal) strongly correlated with the large-scale movement of people (Wang et al., 2020a). Few cases have recorded the cultural diffusion model in the formation of human populations, such as local cultural diffusion with no gene flow events or limited genetic influxes. Wang et al. recently also found two cases of Afanasievo spread into Central Mongolia via the simple cultural diffusion model. To explore the historic demographic model of Chinese Hui, we genotyped 699,537 SNPs in 49 Hui people and 60 geographically close Han people and made one of the most comprehensive whole-genome-based population comparison studies among modern and ancient Eurasian populations. The PCA, ADMIXTURE, pairwise Fst genetic distance, *f*_*3*_, *f*_*4*_, *qpWave/qpAdm*, and *qpGraph*-based admixture graph consistently demonstrated the primary ancestral of Chinese Hui was originated from the Northern East Asian millet farmer populations with little proportion of the ancestry deriving from western Eurasian populations. Here we presented one case of the cultural diffusion model of the Hui people in East Asia.

Previous historic and physical anthropology measurements supported that the culture of Hui people (Muslim) and some of their ancestors spread into China via multiple migration events in different historic times (Yao et al., 2004). Findings from historic documents showed that Silk Road travelers of Persian migrated into Southeast coastal regions of China via Maritime Silk Road, which formed their first stratum. Documents focused on northeast Chinese Hui people showed that they originated from Khorezmians who were transported to as merchants and soldiers by Mongolia empire, which further mixed with local East Asians and other subsequent incoming Central Asians (https://en.wikipedia.org/wiki/Hui_people#cite_note-71). Nowadays, Hui people have their own special cultural beliefs but used the Han Chinese as their language (Leslie and Daniel, 1998;Wang et al., 2019). We found that a small amount of gene flow was derived from the western Eurasian (less than 4∼7 percent) via allele-based *qpAdm* and *qpGraph* analyses. ALDER-based admixture time (about 1000 years ago) further confirmed this historically documented population migration. Thus, the observed western Eurasian ancestry may be the remains of the primary ancestry from West Eurasian. Wang et al. also identified some Y-chromosome lineages enriched in western Eurasian populations, such as western Eurasian related lineages E, F, G, H, I-M, J, L, Q, R, and T (Wang et al., 2019). Dominant East Asian ancestry was identified in this study strongly showed that the main ancestor or ancestry descended from East Asian, suggesting the large-scale genetic assimilation occurred between the original Hui’s ancestor and indigenous East Asians. Y-chromosomal-based this study identified a high proportion of East Eurasian lineages (O1 and O2) and a limited proportion of western Eurasian lineages. Compared to the Southern East Asian, Boshu Hui harbored more ancestry related to Northern East Asian, further supporting Hui’s ancestor mixed with indigenous Northern East Asians. We also observed different demographic histories between Boshui Hui and Guizhou Hui (Wang et al., 2020c), thus, to better understating the genomic history of all geographically diverse Hui people, more genome-wide or sequence data should be collected in the next step.

## Conclusion

We generated genome-wide data in 49 western Hui and 60 Han individuals and merged with all publicly available modern and ancient genomes to conduct one of the most comprehensive population genetic comparisons focused on testing the two contrasting hypotheses of the origin, diversification, migration and admixture of Hui people: East Asian Origin (Cultural diffusion model) and western Eurasian Origin hypothesis (Demic diffusion). We first identified significant genetic differentiation among geographically different Hui or between Hui and adjacent Han populations, suggesting their different demographic structure (**Table 1**). Findings based on the *f*-statistics demonstrated that both Hui and Han possessed stronger Northern East Asian affinity, especially for late Neolithic-to-Iron-Age millet farmers from the middle Yellow River Basin in the Central Plain, supporting their North China Origin. The successfully fitted three-way admixture model with a new-identified small western Eurasian ancestry and dominant northern and southern indigenous East Asian-like ancestry demonstrated that modern East Asian Hui people were mixed via minimal Central Asian with their cultures and massive indigenous Eastern Asian. The mixed populations adopted Muslim culture and ideas, which formed our observed Hui’s gene pool composition, and it consistent with the cultural diffusion model playing a key role in the formation of modern Hui. Besides, the estimated admixture time based on the decay of admixture-introduced linkage disequilibrium revealed the genetic time of introduced western Eurasian ancestry into modern Hui people occurred around 1000 years ago, consistent with the eastern-to-western Eurasian contact during the Tang, Song and Yuan dynasties.

## Acknowledgements

JBY was supported by the Doctoral research fund of North Sichuan Medical College (CBY17-QD13), the Scientific research fund of North Sichuan Medical College and Nanchong Government (19SXHZ0138), and the Scientific research fund of Health Commission of Sichuan Province (18ZD038).

## Data availability statement

All analysis data were presented in the Supplementary Tables. Raw data could be obtained via personal communication with corresponding authors with the population genetic history story only. Which is following the regulations of the Human Genetic Resources Administration of China (HGRAC).

## Author contribution

GH, YL, GC, XZ, and MW conceived the idea for the study. YL, JY, YL, RT, DY, YW, PW, SD, SZ, and HL performed or supervised wet laboratory work. GH, GC, XZ, YL, JY, YL, RT, DY, YW, PW, SD, SZ, and HL analyzed the data. GH, YL, MW and XZ wrote and edited the manuscript.

## Legends of Table

**Table 1**. Differentiated shared alleles between Boshu Hui and Nanchong Han compared to modern and ancient Eurasians estimated via symmetrical *f*_*4*_-statistics in the form *f*_*4*_*(Han_Nanchong, Hui_Boshu; Eurasian, Mbuti)* based on the merged 1240K and Human Origin datasets.

## Legends of Supplementary Tables

**Table S1**. Population-level ancestry composition of the best-fitted thirteen-source ADMIXTURE model.

**Table S2**. The pairwise Fst genetic distance among 237 Eurasian modern and ancient populations.

**Table S3**. The shared genetic drift between Boshu Hui, Nanchong Han and other ancient/modern Eurasians.

**Table S4**. Admixture signature of Nanchong Han population estimated via admixture-*f3*-statistics in the form *f*_*3*_*(Source1, Source2; Han_Nanchong)*.

**Table S5**. Admixture signature of Boshu Hui population estimated via admixture-*f3*-statistics in the form *f*_*3*_*(Source1, Source2; Hui_Boshu)*.

**Table S6**. Differentiated shared alleles between Boshu Hui and Nanchong Han compared to modern and ancient Eurasians estimated via symmetrical *f*_*4*_-statistics in the form *f*_*4*_*(Han_Nanchong, Hui_Boshu; Eurasian, Mbuti)* based on the merged Human Origin dataset.

**Table S7**. Differentiated shared alleles between Boshu Hui and Nanchong Han compared to modern and ancient Eurasians estimated via symmetrical *f*_*4*_-statistics in the form *f*_*4*_*(Han_Nanchong, Hui_Boshu; Eurasian, Mbuti*.*DG)* based on the merged 1240K dataset.

**Table S8**. The results of *f*_*4*_-statistics in the form *f*_*4*_*(Eurasian, Han_Nanchong; Hui_Boshu, Mbuti)* showed genomic affinity between Hui and Han based on the merged Human Origin dataset.

**Table S9**. The results of *f*_*4*_-statistics in the form *f*_*4*_*(Eurasian, Han_Nanchong; Hui_Boshu, Mbuti*.*DG)* showed genomic affinity between Hui and Han based on the merged 1240K dataset.

**Table S10**. Differentiated genomic history between Boshu Hui and Guizhou Hui populations estimated via three *f*_*4*_-statistics

**Table S11**. Symmetrical *f*_*4*-_statistics in the form *f*_*4*_*(Reference population1, reference population2; Hui_Boshu/Han_Nanchong, Mbuti)* to test their genetic affinity between Eurasian reference populations based on the merged 1240K dataset.

**Table S12**. Affinity-*f*_*4*_-statistics in the form *f*_*4*_*(Reference population1, Hui_Boshu/Han_Nanchong; reference population2, Mbuti)* to explore the genetic continuity and admixture between potential ancestral sources and Hui_Boshu/Han_Nanchong based on the merged 1240K dataset.

**Table S13**. Symmetrical *f*_*4*_-statistics in the form *f*_*4*_*(Reference population1, reference population2; Hui_Boshu/Han_Nanchong, Mbuti)* to test their genetic affinity between Eurasian reference populations based on the merged Human Origin dataset.

**Table S14**. Affinity-*f*_*4*_-statistics in the form *f*_*4*_*(Reference population1, Hui_Boshu/Han_Nanchong; reference population2, Mbuti)* to explore the genetic continuity and admixture between potential ancestral sources and Hui_Boshu/Han_Nanchong based on the merged Human Origin dataset.

**Table S15**. qpAdm-based ancestral admixture proportion in Han and Hui via three-way admixture and two-way admixture models.

**Table S16**. Admixture proportion of two-way admixture model for Hui people when used Nanchong Han as the East Asian source.

**Table S17**. Admixture-introduced linkage disequilibrium (ALDER) based admixture time between different eastern and western ancestral sources focused on Uyghur. We used 28 years as the one generation length.

**Table S18**. Admixture-introduced linkage disequilibrium (ALDER) based admixture time between different northern and southern or eastern and western ancestral sources focused on Hui and Han. We used 28 years as the one generation length.

**Table S19**. Haplogroup distribution in Han and Hui populations.

## Legends of Supplementary Figures

**Figure S1**. Geographic position of new collected Hui and Han people from Sichuan Province, Southwest China.

**Figure S2**. Cross-Validation error value of model-based ADMIXTURE analysis.

**Figure S3**. Principal component analysis among Sino-Tibetan speakers and southern East Asians from Austronesian, Austroasiatic, Tai-Kadai and Hmong-Mien language families.

**Figure S4A**. Ancestry compositions of model-based ADMIXTURE analysis with the predefined ancestral sources ranging from two to twenty among modern and ancient Eurasian populations. Left part of the whole picture.

**Figure S4B**. Ancestry compositions of model-based ADMIXTURE analysis with the predefined ancestral sources ranging from two to twenty among modern and ancient Eurasian populations. Right part of the whole picture.

**Figure S5**. Shared genetic drift between the Hui (A), Han (B) and modern Eurasian reference populations estimated via Outgroup-f3-statistics.

**Figure S6**. Excess of sharing alleles between Boshu Hui and modern and ancient Eurasians showed genetic admixture based on the merged 1240K dataset.

**Figure S7**. Excess of sharing alleles between Nanchong Han and modern and ancient Eurasians.

**Figure S8**. Results of f4-statistics showed genomic relationship inferred from f4(Reference population1, reference population2; Hui_Boshu, Mbuti) based on the merged Human Origin dataset.

**Figure S9**. Results of f4-statistics showed genomic relationship inferred from f4(Reference population1, reference population2; Han_Nanchong, Mbuti) based on the merged Human Origin dataset.

**Figure S10**. Excess of sharing alleles between Nanchong Han and modern and ancient Eurasians based on the merged Human Origin dataset.

**Figure S11**. qpGraph-based admixture graph showed the western Eurasian gene flow event related to Loschbour (A) and Kazakhstan_Andronovo (B) in Boshu Hui.

**Figure S12**. qpGraph-based admixture graph showed the western Eurasian gene flow event related to Russia_Sintashta_MLBA (A) and Russia_Alan (B) in Boshu Hui.

**Figure S13**. qpGraph-based admixture graph showed the western Eurasian gene flow event related to Sardinian (A) and French (B) in Boshu Hui.

**Figure S14**. Admixture-introduced linkage disequilibrium (ALDER) based admixture time between different northern and southern or eastern and western ancestral sources.

## References

Alexander, D.H., Novembre, J., and Lange, K. (2009). Fast model-based estimation of ancestry in unrelated individuals. Genome Res 19, 1655–1664.

Allentoft, M.E., Sikora, M., Sjogren, K.G., Rasmussen, S., Rasmussen, M., Stenderup, J., Damgaard, P.B., Schroeder, H., Ahlstrom, T., Vinner, L., Malaspinas, A.S., Margaryan, A., Higham, T., Chivall, D., Lynnerup, N., Harvig, L., Baron, J., Della Casa, P., Dabrowski, P., Duffy, P.R., Ebel, A.V., Epimakhov, A., Frei, K., Furmanek, M., Gralak, T., Gromov, A., Gronkiewicz, S., Grupe, G., Hajdu, T., Jarysz, R., Khartanovich, V., Khokhlov, A., Kiss, V., Kolar, J., Kriiska, A., Lasak, I., Longhi, C., Mcglynn, G., Merkevicius, A., Merkyte, I., Metspalu, M., Mkrtchyan, R., Moiseyev, V., Paja, L., Palfi, G., Pokutta, D., Pospieszny, L., Price, T.D., Saag, L., Sablin, M., Shishlina, N., Smrcka, V., Soenov, V.I., Szeverenyi, V., Toth, G., Trifanova, S.V., Varul, L., Vicze, M., Yepiskoposyan, L., Zhitenev, V., Orlando, L., Sicheritz-Ponten, T., Brunak, S., Nielsen, R., Kristiansen, K., and Willerslev, E. (2015). Population genomics of Bronze Age Eurasia. Nature 522, 167–172.

Association, W.M. (2001). World Medical Association Declaration of Helsinki. Ethical principles for medical research involving human subjects. Bulletin of the World Health Organization 79, 373.

Chang, C.C., Chow, C.C., Tellier, L.C., Vattikuti, S., Purcell, S.M., and Lee, J.J. (2015). Second-generation PLINK: rising to the challenge of larger and richer datasets. Gigascience 4, 7.

Damgaard, P.B., Marchi, N., Rasmussen, S., Peyrot, M., Renaud, G., Korneliussen, T., Moreno-Mayar, J.V., Pedersen, M.W., Goldberg, A., Usmanova, E., Baimukhanov, N., Loman, V., Hedeager, L., Pedersen, A.G., Nielsen, K., Afanasiev, G., Akmatov, K., Aldashev, A., Alpaslan, A., Baimbetov, G., Bazaliiskii, V.I., Beisenov, A., Boldbaatar, B., Boldgiv, B., Dorzhu, C., Ellingvag, S., Erdenebaatar, D., Dajani, R., Dmitriev, E., Evdokimov, V., Frei, K.M., Gromov, A., Goryachev, A., Hakonarson, H., Hegay, T., Khachatryan, Z., Khaskhanov, R., Kitov, E., Kolbina, A., Kubatbek, T., Kukushkin, A., Kukushkin, I., Lau, N., Margaryan, A., Merkyte, I., Mertz, I.V., Mertz, V.K., Mijiddorj, E., Moiyesev, V., Mukhtarova, G., Nurmukhanbetov, B., Orozbekova, Z., Panyushkina, I., Pieta, K., Smrcka, V., Shevnina, I., Logvin, A., Sjogren, K.G., Stolcova, T., Taravella, A.M., Tashbaeva, K., Tkachev, A., Tulegenov, T., Voyakin, D., Yepiskoposyan, L., Undrakhbold, S., Varfolomeev, V., Weber, A., Wilson Sayres, M.A., Kradin, N., Allentoft, M.E., Orlando, L., Nielsen, R., Sikora, M., Heyer, E., Kristiansen, K., and Willerslev, E. (2018). 137 ancient human genomes from across the Eurasian steppes. Nature 557, 369–374.

De Barros Damgaard, P., Martiniano, R., Kamm, J., Moreno-Mayar, J.V., Kroonen, G., Peyrot, M., Barjamovic, G., Rasmussen, S., Zacho, C., Baimukhanov, N., Zaibert, V., Merz, V., Biddanda, A., Merz, I., Loman, V., Evdokimov, V., Usmanova, E., Hemphill, B., Seguin-Orlando, A., Yediay, F.E., Ullah, I., Sjögren, K.-G., Iversen, K.H., Choin, J., De La Fuente, C., Ilardo, M., Schroeder, H., Moiseyev, V., Gromov, A., Polyakov, A., Omura, S., Senyurt, S.Y., Ahmad, H., Mckenzie, C., Margaryan, A., Hameed, A., Samad, A., Gul, N., Khokhar, M.H., Goriunova, O.I., Bazaliiskii, V.I., Novembre, J., Weber, A.W., Orlando, L., Allentoft, M.E., Nielsen, R., Kristiansen, K., Sikora, M., Outram, A.K., Durbin, R., and Willerslev, E. (2018). The first horse herders and the impact of early Bronze Age steppe expansions into Asia. Science 360, eaar7711.

Dong, G., Du, L., and Wei, W. (2020). The impact of early trans-Eurasian exchange on animal utilization in northern China during 5000–2500 BP. The Holocene, 0959683620941169.

Green, R.E., Krause, J., Briggs, A.W., Maricic, T., Stenzel, U., Kircher, M., Patterson, N., Li, H., Zhai, W., Fritz, M.H., Hansen, N.F., Durand, E.Y., Malaspinas, A.S., Jensen, J.D., Marques-Bonet, T., Alkan, C., Prufer, K., Meyer, M., Burbano, H.A., Good, J.M., Schultz, R., Aximu-Petri, A., Butthof, A., Hober, B., Hoffner, B., Siegemund, M., Weihmann, A., Nusbaum, C., Lander, E.S., Russ, C., Novod, N., Affourtit, J., Egholm, M., Verna, C., Rudan, P., Brajkovic, D., Kucan, Z., Gusic, I., Doronichev, V.B., Golovanova, L.V., Lalueza-Fox, C., De La Rasilla, M., Fortea, J., Rosas, A., Schmitz, R.W., Johnson, P.L.F., Eichler, E.E., Falush, D., Birney, E., Mullikin, J.C., Slatkin, M., Nielsen, R., Kelso, J., Lachmann, M., Reich, D., and Paabo, S. (2010). A draft sequence of the Neandertal genome. Science 328, 710–722.

He, G., Wang, Z., Wang, M., Luo, T., Liu, J., Zhou, Y., Gao, B., and Hou, Y. (2018). Forensic ancestry analysis in two Chinese minority populations using massively parallel sequencing of 165 ancestry-informative SNPs. Electrophoresis 39, 2732–2742.

Jeong, C., Wang, K., Wilkin, S., Taylor, W.T.T., Miller, B.K., Bemmann, J.H., Stahl, R., Chiovelli, C., Knolle, F., Ulziibayar, S., Khatanbaatar, D., Erdenebaatar, D., Erdenebat, U., Ochir, A., Ankhsanaa, G., Vanchigdash, C., Ochir, B., Munkhbayar, C., Tumen, D., Kovalev, A., Kradin, N., Bazarov, B.A., Miyagashev, D.A., Konovalov, P.B., Zhambaltarova, E., Miller, A.V., Haak, W., Schiffels, S., Krause, J., Boivin, N., Erdene, M., Hendy, J., and Warinner, C. (2020). A Dynamic 6,000-Year Genetic History of Eurasia’s Eastern Steppe. Cell 183, 890–904 e829.

Kumar, S., Stecher, G., and Tamura, K. (2016). MEGA7: Molecular Evolutionary Genetics Analysis Version 7.0 for Bigger Datasets. Mol Biol Evol 33, 1870–1874.

Lazaridis, I., Patterson, N., Mittnik, A., Renaud, G., Mallick, S., Kirsanow, K., Sudmant, P.H., Schraiber, J.G., Castellano, S., Lipson, M., Berger, B., Economou, C., Bollongino, R., Fu, Q., Bos, K.I., Nordenfelt, S., Li, H., De Filippo, C., Prufer, K., Sawyer, S., Posth, C., Haak, W., Hallgren, F., Fornander, E., Rohland, N., Delsate, D., Francken, M., Guinet, J.M., Wahl, J., Ayodo, G., Babiker, H.A., Bailliet, G., Balanovska, E., Balanovsky, O., Barrantes, R., Bedoya, G., Ben-Ami, H., Bene, J., Berrada, F., Bravi, C.M., Brisighelli, F., Busby, G.B., Cali, F., Churnosov, M., Cole, D.E., Corach, D., Damba, L., Van Driem, G., Dryomov, S., Dugoujon, J.M., Fedorova, S.A., Gallego Romero, I., Gubina, M., Hammer, M., Henn, B.M., Hervig, T., Hodoglugil, U., Jha, A.R., Karachanak-Yankova, S., Khusainova, R., Khusnutdinova, E., Kittles, R., Kivisild, T., Klitz, W., Kucinskas, V., Kushniarevich, A., Laredj, L., Litvinov, S., Loukidis, T., Mahley, R.W., Melegh, B., Metspalu, E., Molina, J., Mountain, J., Nakkalajarvi, K., Nesheva, D., Nyambo, T., Osipova, L., Parik, J., Platonov, F., Posukh, O., Romano, V., Rothhammer, F., Rudan, I., Ruizbakiev, R., Sahakyan, H., Sajantila, A., Salas, A., Starikovskaya, E.B., Tarekegn, A., Toncheva, D., Turdikulova, S., Uktveryte, I., Utevska, O., Vasquez, R., Villena, M., Voevoda, M., Winkler, C.A., Yepiskoposyan, L., Zalloua, P., et al. (2014). Ancient human genomes suggest three ancestral populations for present-day Europeans. Nature 513, 409–413.

Leipe, C., Long, T., Sergusheva, E.A., Wagner, M., and Tarasov, P.E. (2019). Discontinuous spread of millet agriculture in eastern Asia and prehistoric population dynamics. Sci Adv 5, eaax6225.

Leslie, and Daniel, D. (1998). The Integration of Religious Minorities in China: The Case of Chinese Muslims.

Loh, P.R., Lipson, M., Patterson, N., Moorjani, P., Pickrell, J.K., Reich, D., and Berger, B. (2013). Inferring admixture histories of human populations using linkage disequilibrium. Genetics 193, 1233–1254.

Narasimhan, V.M., Patterson, N., Moorjani, P., Rohland, N., Bernardos, R., Mallick, S., Lazaridis, I., Nakatsuka, N., Olalde, I., Lipson, M., Kim, A.M., Olivieri, L.M., Coppa, A., Vidale, M., Mallory, J., Moiseyev, V., Kitov, E., Monge, J., Adamski, N., Alex, N., Broomandkhoshbacht, N., Candilio, F., Callan, K., Cheronet, O., Culleton, B.J., Ferry, M., Fernandes, D., Freilich, S., Gamarra, B., Gaudio, D., Hajdinjak, M., Harney, E., Harper, T.K., Keating, D., Lawson, A.M., Mah, M., Mandl, K., Michel, M., Novak, M., Oppenheimer, J., Rai, N., Sirak, K., Slon, V., Stewardson, K., Zalzala, F., Zhang, Z., Akhatov, G., Bagashev, A.N., Bagnera, A., Baitanayev, B., Bendezu-Sarmiento, J., Bissembaev, A.A., Bonora, G.L., Chargynov, T.T., Chikisheva, T., Dashkovskiy, P.K., Derevianko, A., Dobes, M., Douka, K., Dubova, N., Duisengali, M.N., Enshin, D., Epimakhov, A., Fribus, A.V., Fuller, D., Goryachev, A., Gromov, A., Grushin, S.P., Hanks, B., Judd, M., Kazizov, E., Khokhlov, A., Krygin, A.P., Kupriyanova, E., Kuznetsov, P., Luiselli, D., Maksudov, F., Mamedov, A.M., Mamirov, T.B., Meiklejohn, C., Merrett, D.C., Micheli, R., Mochalov, O., Mustafokulov, S., Nayak, A., Pettener, D., Potts, R., Razhev, D., Rykun, M., Sarno, S., Savenkova, T.M., Sikhymbaeva, K., Slepchenko, S.M., Soltobaev, O.A., Stepanova, N., Svyatko, S., Tabaldiev, K., Teschler-Nicola, M., Tishkin, A.A., Tkachev, V.V., et al. (2019). The formation of human populations in South and Central Asia. Science 365, eaat7487.

Ning, C., Li, T., Wang, K., Zhang, F., Li, T., Wu, X., Gao, S., Zhang, Q., Zhang, H., Hudson, M.J., Dong, G., Wu, S., Fang, Y., Liu, C., Feng, C., Li, W., Han, T., Li, R., Wei, J., Zhu, Y., Zhou, Y., Wang, C.C., Fan, S., Xiong, Z., Sun, Z., Ye, M., Sun, L., Wu, X., Liang, F., Cao, Y., Wei, X., Zhu, H., Zhou, H., Krause, J., Robbeets, M., Jeong, C., and Cui, Y. (2020). Ancient genomes from northern China suggest links between subsistence changes and human migration. Nat Commun 11, 2700.

Ning, C., Wang, C.C., Gao, S., Yang, Y., Zhang, X., Wu, X., Zhang, F., Nie, Z., Tang, Y., Robbeets, M., Ma, J., Krause, J., and Cui, Y. (2019). Ancient Genomes Reveal Yamnaya-Related Ancestry and a Potential Source of Indo-European Speakers in Iron Age Tianshan. Curr Biol 29, 2526–2532 e2524.

Patin, E., Lopez, M., Grollemund, R., Verdu, P., Harmant, C., Quach, H., Laval, G., Perry, G.H., Barreiro, L.B., Froment, A., Heyer, E., Massougbodji, A., Fortes-Lima, C., Migot-Nabias, F., Bellis, G., Dugoujon, J.M., Pereira, J.B., Fernandes, V., Pereira, L., Van Der Veen, L., Mouguiama-Daouda, P., Bustamante, C.D., Hombert, J.M., and Quintana-Murci, L. (2017). Dispersals and genetic adaptation of Bantu-speaking populations in Africa and North America. Science 356, 543–546.

Patterson, N., Moorjani, P., Luo, Y., Mallick, S., Rohland, N., Zhan, Y., Genschoreck, T., Webster, T., and Reich, D. (2012). Ancient admixture in human history. Genetics 192, 1065–1093.

Pickrell, J.K., and Pritchard, J.K. (2012). Inference of population splits and mixtures from genome-wide allele frequency data. PLoS Genet 8, e1002967.

Raghavan, M., Steinrucken, M., Harris, K., Schiffels, S., Rasmussen, S., Degiorgio, M., Albrechtsen, A., Valdiosera, C., Avila-Arcos, M.C., Malaspinas, A.S., Eriksson, A., Moltke, I., Metspalu, M., Homburger, J.R., Wall, J., Cornejo, O.E., Moreno-Mayar, J.V., Korneliussen, T.S., Pierre, T., Rasmussen, M., Campos, P.F., De Barros Damgaard, P., Allentoft, M.E., Lindo, J., Metspalu, E., Rodriguez-Varela, R., Mansilla, J., Henrickson, C., Seguin-Orlando, A., Malmstrom, H., Stafford, T., Jr., Shringarpure, S.S., Moreno-Estrada, A., Karmin, M., Tambets, K., Bergstrom, A., Xue, Y., Warmuth, V., Friend, A.D., Singarayer, J., Valdes, P., Balloux, F., Leboreiro, I., Vera, J.L., Rangel-Villalobos, H., Pettener, D., Luiselli, D., Davis, L.G., Heyer, E., Zollikofer, C.P.E., Ponce De Leon, M.S., Smith, C.I., Grimes, V., Pike, K.A., Deal, M., Fuller, B.T., Arriaza, B., Standen, V., Luz, M.F., Ricaut, F., Guidon, N., Osipova, L., Voevoda, M.I., Posukh, O.L., Balanovsky, O., Lavryashina, M., Bogunov, Y., Khusnutdinova, E., Gubina, M., Balanovska, E., Fedorova, S., Litvinov, S., Malyarchuk, B., Derenko, M., Mosher, M.J., Archer, D., Cybulski, J., Petzelt, B., Mitchell, J., Worl, R., Norman, P.J., Parham, P., Kemp, B.M., Kivisild, T., Tyler-Smith, C., Sandhu, M.S., Crawford, M., Villems, R., Smith, D.G., Waters, M.R., Goebel, T., Johnson, J.R., Malhi, R.S., Jakobsson, M., Meltzer, D.J., Manica, A., Durbin, R., Bustamante, C.D., Song, Y.S., Nielsen, R., et al. (2015). POPULATION GENETICS. Genomic evidence for the Pleistocene and recent population history of Native Americans. Science 349, aab3884.

Reich, D., Green, R.E., Kircher, M., Krause, J., Patterson, N., Durand, E.Y., Viola, B., Briggs, A.W., Stenzel, U., Johnson, P.L., Maricic, T., Good, J.M., Marques-Bonet, T., Alkan, C., Fu, Q., Mallick, S., Li, H., Meyer, M., Eichler, E.E., Stoneking, M., Richards, M., Talamo, S., Shunkov, M.V., Derevianko, A.P., Hublin, J.J., Kelso, J., Slatkin, M., and Paabo, S. (2010). Genetic history of an archaic hominin group from Denisova Cave in Siberia. Nature 468, 1053–1060.

Sikora, M., Pitulko, V.V., Sousa, V.C., Allentoft, M.E., Vinner, L., Rasmussen, S., Margaryan, A., De Barros Damgaard, P., De La Fuente, C., Renaud, G., Yang, M.A., Fu, Q., Dupanloup, I., Giampoudakis, K., Nogues-Bravo, D., Rahbek, C., Kroonen, G., Peyrot, M., Mccoll, H., Vasilyev, S.V., Veselovskaya, E., Gerasimova, M., Pavlova, E.Y., Chasnyk, V.G., Nikolskiy, P.A., Gromov, A.V., Khartanovich, V.I., Moiseyev, V., Grebenyuk, P.S., Fedorchenko, A.Y., Lebedintsev, A.I., Slobodin, S.B., Malyarchuk, B.A., Martiniano, R., Meldgaard, M., Arppe, L., Palo, J.U., Sundell, T., Mannermaa, K., Putkonen, M., Alexandersen, V., Primeau, C., Baimukhanov, N., Malhi, R.S., Sjogren, K.G., Kristiansen, K., Wessman, A., Sajantila, A., Lahr, M.M., Durbin, R., Nielsen, R., Meltzer, D.J., Excoffier, L., and Willerslev, E. (2019). The population history of northeastern Siberia since the Pleistocene. Nature 570, 182–188.

Wang, C.-C., and Robbeets, M. (2020). The homeland of Proto-Tungusic inferred from contemporary words and ancient genomes. Evolutionary Human Sciences 2.

Wang, C.-C., Yeh, H.-Y., Popov, A.N., Zhang, H.-Q., Matsumura, H., Sirak, K., Cheronet, O., Kovalev, A., Rohland, N., Kim, A.M., Bernardos, R., Tumen, D., Zhao, J., Liu, Y.-C., Liu, J.-Y., Mah, M., Mallick, S., Wang, K., Zhang, Z., Adamski, N., Broomandkhoshbacht, N., Callan, K., Culleton, B.J., Eccles, L., Lawson, A.M., Michel, M., Oppenheimer, J., Stewardson, K., Wen, S., Yan, S., Zalzala, F., Chuang, R., Huang, C.-J., Shiung, C.-C., Nikitin, Y.G., Tabarev, A.V., Tishkin, A.A., Lin, S., Sun, Z.-Y., Wu, X.-M., Yang, T.-L., Hu, X., Chen, L., Du, H., Bayarsaikhan, J., Mijiddorj, E., Erdenebaatar, D., Iderkhangai, T.-O., Myagmar, E., Kanzawa-Kiriyama, H., Nishino, M., Shinoda, K.-I., Shubina, O.A., Guo, J., Deng, Q., Kang, L., Li, D., Li, D., Lin, R., Cai, W., Shrestha, R., Wang, L.-X., Wei, L., Xie, G., Yao, H., Zhang, M., He, G., Yang, X., Hu, R., Robbeets, M., Schiffels, S., Kennett, D.J., Jin, L., Li, H., Krause, J., Pinhasi, R., and Reich, D. (2020a). The Genomic Formation of Human Populations in East Asia. bioRxiv, 2020.2003.2025.004606.

Wang, C.C., Lu, Y., Kang, L., Ding, H., Yan, S., Guo, J., Zhang, Q., Wen, S.Q., Wang, L.X., Zhang, M., Tong, X., Huang, X., Nie, S., Deng, Q., Zhu, B., Jin, L., and Li, H. (2019). The massive assimilation of indigenous East Asian populations in the origin of Muslim Hui people inferred from paternal Y chromosome. Am J Phys Anthropol 169, 341–347.

Wang, M., Zou, X., Ye, H.-Y., Wang, Z., Liu, Y., Liu, J., Wang, F., Yao, H., Chen, P., Tao, R., Wang, S., Wei, L.-H., Tang, R., Wang, C.-C., and He, G. (2020b). Peopling of Tibet Plateau and multiple waves of admixture of Tibetans inferred from both modern and ancient genome-wide data. bioRxiv, 2020.2007.2003.185884.

Wang, Q., Zhao, J., Ren, Z., Sun, J., He, G., Guo, J., Zhang, H., Ji, J., Liu, Y., Yang, M., Yang, X., Chen, J., Zhu, K., Wang, R., Li, Y., Chen, G., Huang, J., and Wang, C.-C. (2020c). Male-dominated migration and massive assimilation of indigenous East Asians in the formation of Muslim Hui people in southwest China. Frontiers in Genetics.

Weir, B.S., and Cockerham, C.C. (1984). Estimating F-Statistics for the Analysis of Population Structure. Evolution 38, 1358–1370.

Weissensteiner, H., Pacher, D., Kloss-Brandstätter, A., Forer, L., Specht, G., Bandelt, H.-J., Kronenberg, F., Salas, A., and Schönherr, S. (2016). HaploGrep 2: mitochondrial haplogroup classification in the era of high-throughput sequencing. Nucleic Acids Research 44, W58–W63.

Wen, B., Li, H., Lu, D., Song, X., Zhang, F., He, Y., Li, F., Gao, Y., Mao, X., Zhang, L., Qian, J., Tan, J., Jin, J., Huang, W., Deka, R., Su, B., Chakraborty, R., and Jin, L. (2004). Genetic evidence supports demic diffusion of Han culture. Nature 431, 302–305.

Yang, M.A., Fan, X., Sun, B., Chen, C., Lang, J., Ko, Y.C., Tsang, C.H., Chiu, H., Wang, T., Bao, Q., Wu, X., Hajdinjak, M., Ko, A.M., Ding, M., Cao, P., Yang, R., Liu, F., Nickel, B., Dai, Q., Feng, X., Zhang, L., Sun, C., Ning, C., Zeng, W., Zhao, Y., Zhang, M., Gao, X., Cui, Y., Reich, D., Stoneking, M., and Fu, Q. (2020). Ancient DNA indicates human population shifts and admixture in northern and southern China. Science 369, 282–288.

Yao, H.B., Wang, C.C., Tao, X., Shang, L., Wen, S.Q., Zhu, B., Kang, L., Jin, L., and Li, H. (2016). Genetic evidence for an East Asian origin of Chinese Muslim populations Dongxiang and Hui. Sci Rep 6, 38656.

Yao, Y.G., Kong, Q.P., Wang, C.Y., Zhu, C.L., and Zhang, Y.P. (2004). Different matrilineal contributions to genetic structure of ethnic groups in the silk road region in china. Mol Biol Evol 21, 2265–2280.

Yunusbayev, B., Metspalu, M., Metspalu, E., Valeev, A., Litvinov, S., Valiev, R., Akhmetova, V., Balanovska, E., Balanovsky, O., Turdikulova, S., Dalimova, D., Nymadawa, P., Bahmanimehr, A., Sahakyan, H., Tambets, K., Fedorova, S., Barashkov, N., Khidiyatova, I., Mihailov, E., Khusainova, R., Damba, L., Derenko, M., Malyarchuk, B., Osipova, L., Voevoda, M., Yepiskoposyan, L., Kivisild, T., Khusnutdinova, E., and Villems, R. (2015). The genetic legacy of the expansion of Turkic-speaking nomads across Eurasia. PLoS Genet 11, e1005068.

Zhou, B., Wen, S., Sun, H., Zhang, H., and Shi, R. (2020). Genetic affinity between Ningxia Hui and eastern Asian populations revealed by a set of InDel loci. R Soc Open Sci 7, 190358.

Zou, X., Wang, Z., He, G., Wang, M., Liu, J., Wang, S., Ye, Z., Wang, F., and Hou, Y. (2020). Genetic variation and population structure analysis of Chinese Wuzhong Hui population using 30 Indels. Ann Hum Biol 47, 300–303.

