## Supplementary Figure S1∼18 for "Significant East Asian affinity of Chinese Hui genomic structure suggesting their predominant cultural diffusion model in the genetic formation process"

[1]School of Basic Medical Sciences, North Sichuan Medical College, Nanchong, Sichuan, 637100, China
[2]AnLan AI, Shenzhen, China
[3]Department of Forensic Medicine, College of Basic Medicine, Chongqing Medical University, Chongqing, 400331, China
[4]College of Medical Information, Chongqing Medical University, Chongqing, 400331, China
[5]School of Medical Imaging, North Sichuan Medical College, Nanchong, Sichuan, 637000, China
[6]Hunan Key Lab of Bioinformatics, School of Computer Science and Engineering, Central South University, Changsha, 410075, China
[7]Institute of Forensic Medicine, West China School of Basic Science and Forensic Medicine, Sichuan University, Chengdu, 610065, China
[8]Department of Anthropology and Ethnology, Institute of Anthropology, National Institute for Data Science in Health and Medicine, and School of Life Sciences, Xiamen University, Xiamen, 361005, China
*These authors contributed equally to this work and should be considered co-first authors.
#Corresponding author
**Gang Chen**
**Affiliation**: Hunan Key Lab of Bioinformatics, School of Computer Science and Engineering, Central South University, Changsha, 410075, China

**Xing Zou**
**Affiliation**: Institute of Forensic Medicine, West China School of Basic Science and Forensic Medicine, Sichuan University
**Mengge Wang**
**Affiliation**: Institute of Forensic Medicine, West China School of Basic Science and Forensic Medicine, Sichuan University
**Guanglin He**
**Affiliation**: Department of Anthropology and Ethnology, Institute of Anthropology, National Institute for Data Science in Health and Medicine, Xiamen University, Xiamen, China.

### Supplementary Tables S1-18

Figure S1. Geographic position of new collected Hui and Han people from Sichuan Province, Southwest China. 3
Figure S2. Cross-Validation error value of model-based ADMIXTURE analysis. 3
Figure S3. Principal component analysis among Sino-Tibetan speakers and southern East Asians from Austronesian, Austroasiatic, Tai-Kadai and Hmong-Mien language families. 4
Figure S4A. Ancestry compositions of model-based ADMIXTURE analysis with the predefined ancestral sources ranging from two to twenty among modern and ancient Eurasian populations. Left part of the whole picture. 5
Figure S4B. Ancestry compositions of model-based ADMIXTURE analysis with the predefined ancestral sources ranging from two to twenty among modern and ancient Eurasian populations. Right part of the whole picture. 6
Figure S5. Shared genetic drift between the Hui (A), Han (B) and modern Eurasian reference populations estimated via Outgroup-$f_3$-statistics. 7
Figure S6. Excess of sharing alleles between Boshu Hui and modern and ancient Eurasians showed genetic admixture based on the merged 1240K dataset. 8
Figure S7. Excess of sharing alleles between Nanchong Han and modern and ancient Eurasians. 9
Figure S8. Results of $f_4$-statistics showed genomic relationship inferred from $f_4$*(Reference population1, reference population2; Hui_Boshu, Mbuti)* based on the merged Human Origin dataset. 10
Figure S9. Results of $f_4$-statistics showed genomic relationship inferred from $f_4$*(Reference population1, reference population2; Han_Nanchong, Mbuti)* based on the merged Human Origin dataset. 11
Figure S10. Excess of sharing alleles between Nanchong Han and modern and ancient Eurasians based on the merged Human Origin dataset. 12
Figure S11. *qpGraph*-based admixture graph showed the western Eurasian gene flow event related to Kalmykia Yamnaya into Boshu Hui. 13
Figure S12. *qpGraph*-based admixture graph showed the western Eurasian gene flow event related to Russia_Sintashta_MLBA into Boshu Hui. 14
Figure S13. *qpGraph*-based admixture graph showed the western Eurasian gene flow event related to Russia Alan into Boshu Hui. 15
Figure S14. *qpGraph*-based admixture graph showed the western Eurasian gene flow event related to Moldova Scythian into Boshu Hui. 16
Figure S15. *qpGraph*-based admixture graph showed the western Eurasian gene flow event related to French into Boshu Hui. 17
Figure S16. *qpGraph*-based admixture graph showed the western Eurasian gene flow event related to Sardinian into Boshu Hui. 18
Figure S17. *qpGraph*-based admixture graph showed the western Eurasian gene flow event related to Spanish into Boshu Hui. 19
Figure S18. Admixture-introduced linkage disequilibrium (ALDER) based admixture time between different northern and southern or eastern and western ancestral sources. 20

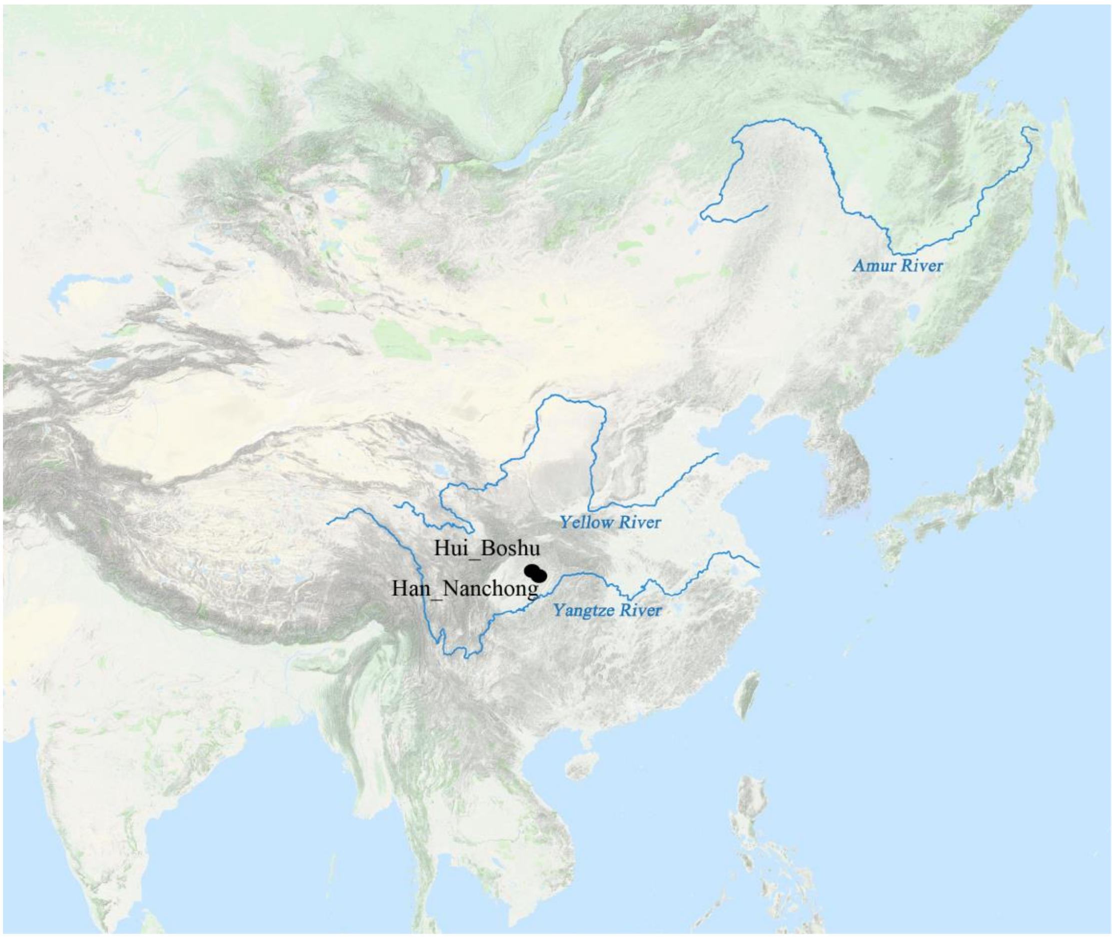


**Figure S1. Geographic position of new collected Hui and Han people from Sichuan Province, Southwest China.**

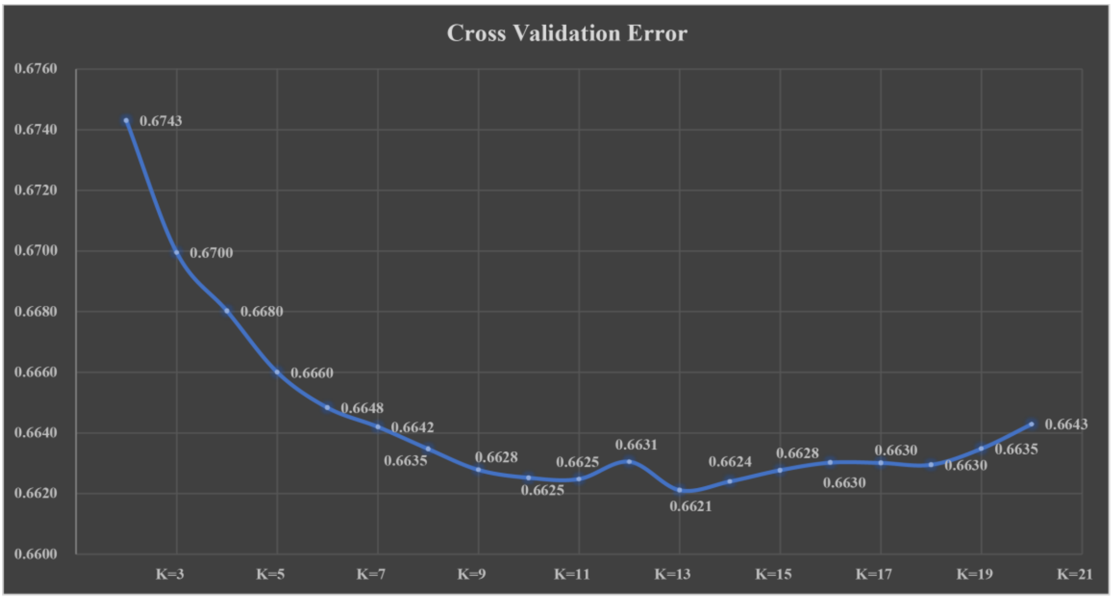


**Figure S2. Cross-Validation error value of model-based ADMIXTURE analysis.**

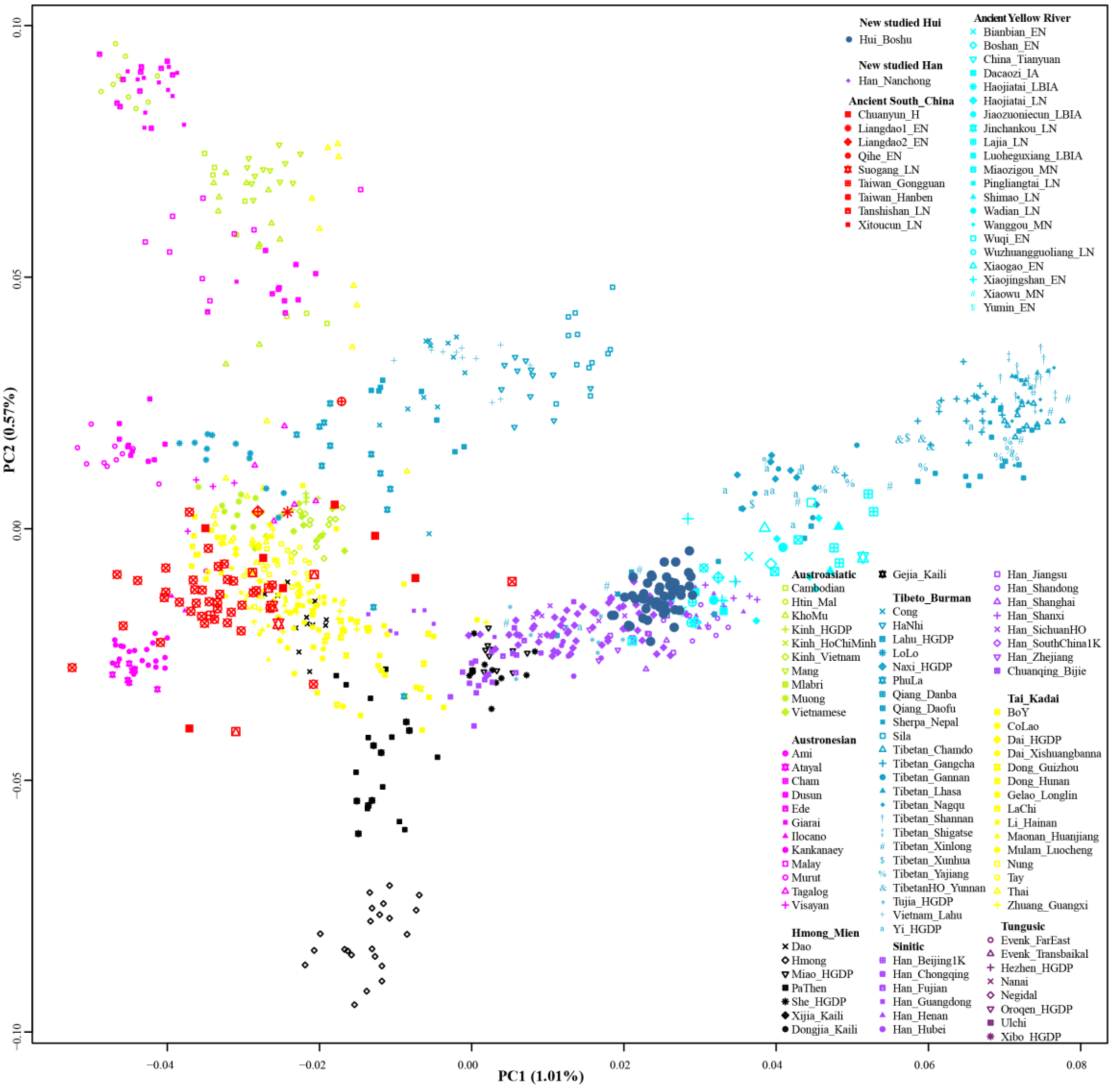


**Figure S3. Principal component analysis among Sino-Tibetan speakers and southern East Asians from Austronesian, Austroasiatic, Tai-Kadai and Hmong-Mien language families.**

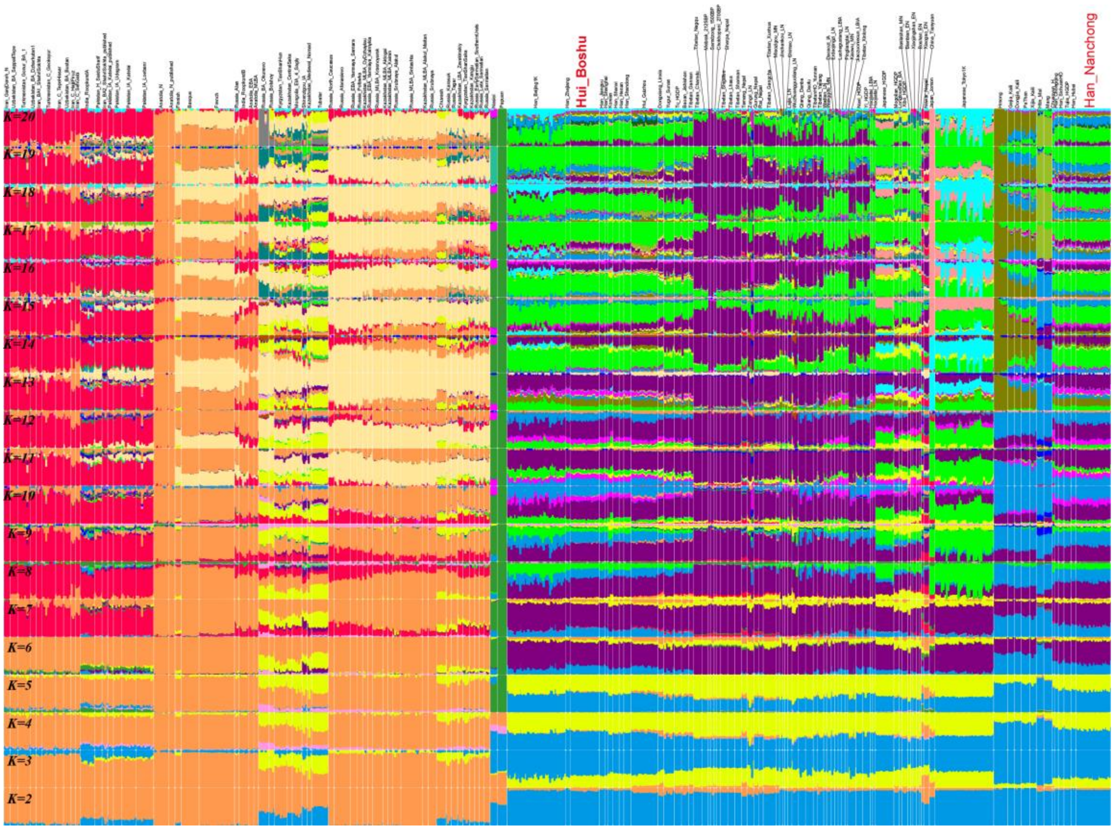


**Figure S4A. Ancestry compositions of model-based ADMIXTURE analysis with the predefined ancestral sources ranging from two to twenty among modern and ancient Eurasian populations. Left part of the whole picture.**

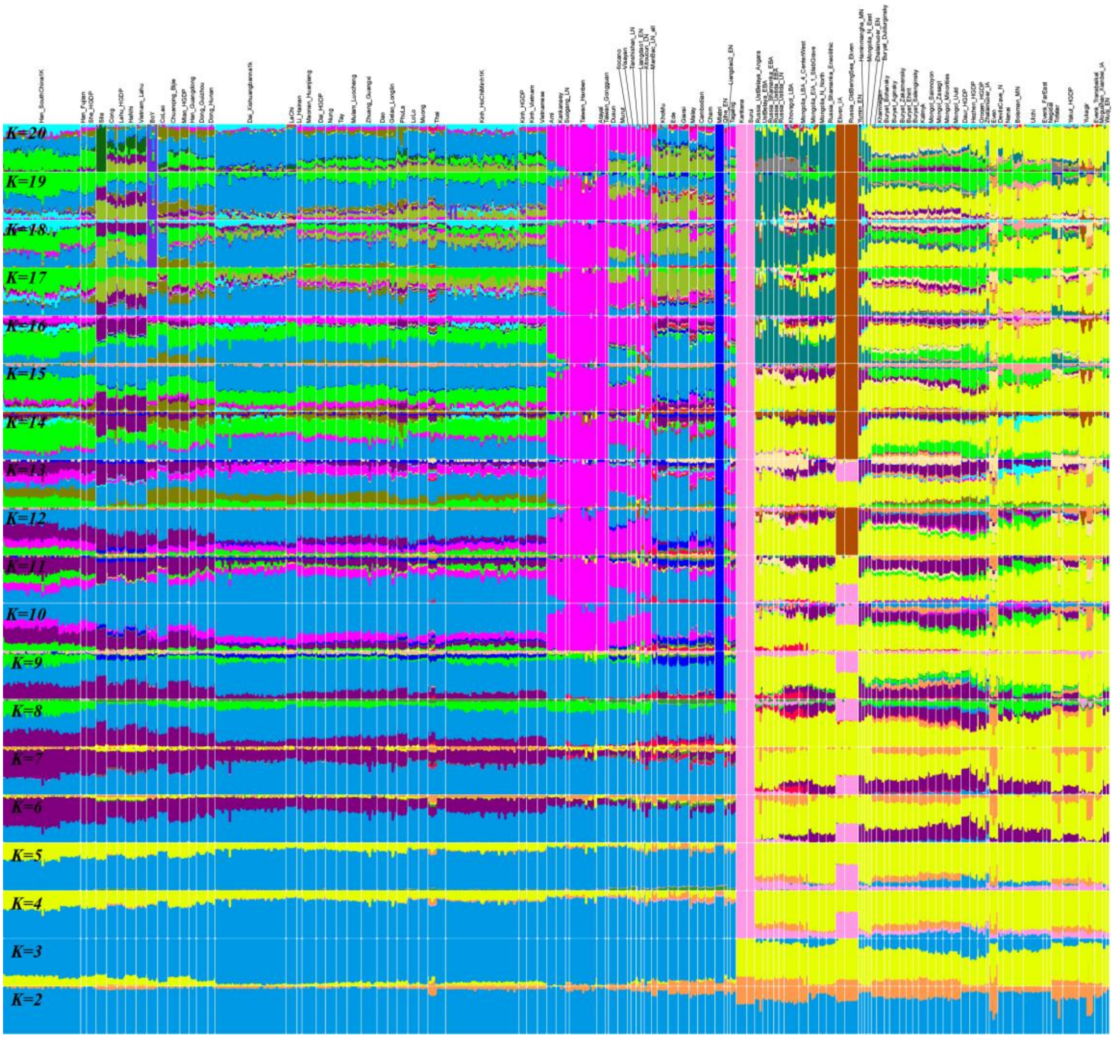


**Figure S4B. Ancestry compositions of model-based ADMIXTURE analysis with the predefined ancestral sources ranging from two to twenty among modern and ancient Eurasian populations. Right part of the whole picture.**

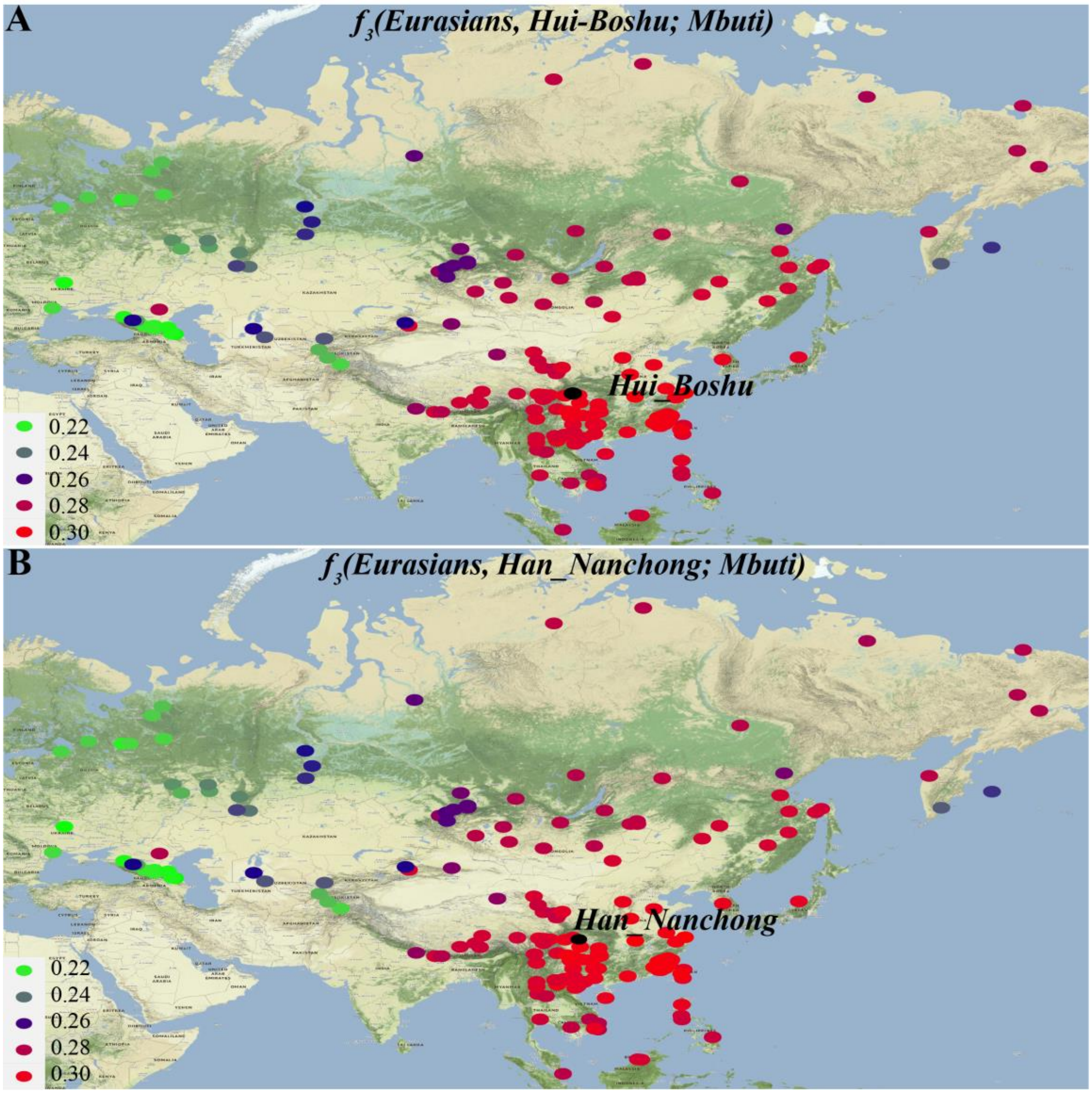


**Figure S5. Shared genetic drift between the Hui (A), Han (B) and modern Eurasian reference populations estimated via Outgroup-$f_3$-statistics.**

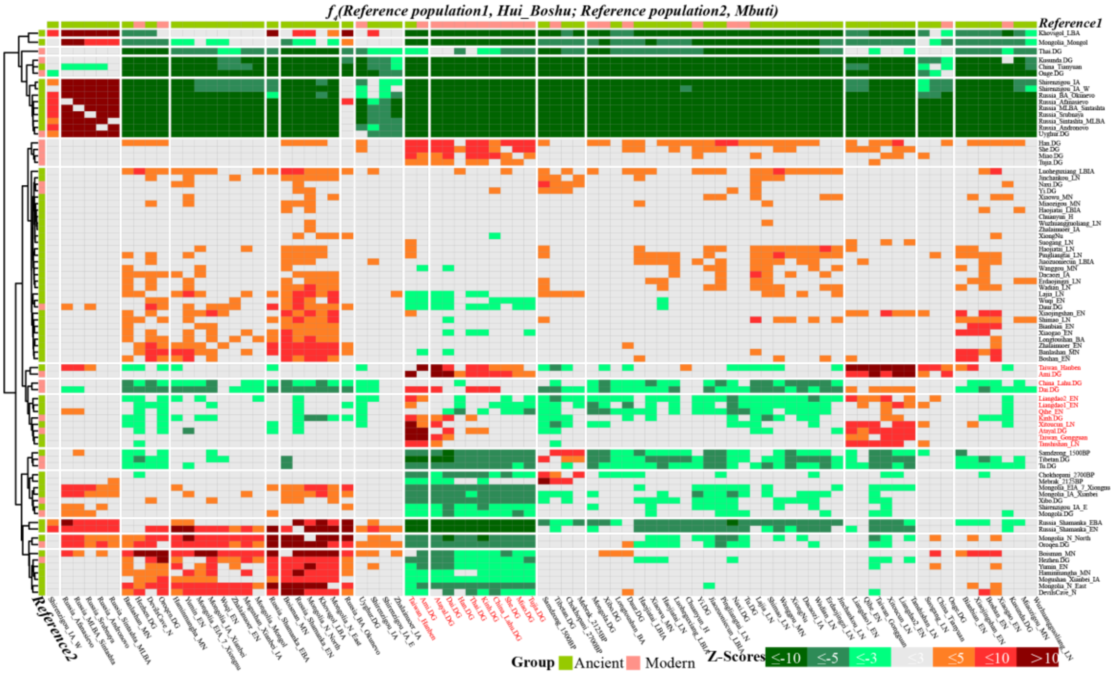


**Figure S6. Excess of sharing alleles between Boshu Hui and modern and ancient Eurasians showed genetic admixture based on the merged 1240K dataset.**

Affinity-$f_4$-statistics in the form $f_4$*(Reference population1, Hui_Boshu; reference population2, Mbuti)* to explore the genetic continuity and admixture between potential ancestral sources and Boshu Hui based on the merged 1240K dataset. The red color denoted the third population (Bottom population lists) shared more derived alleles with the first population (Right population lists) compared with the second one. And the green color denoted the third population shared more alleles with the second one relative to the first one.

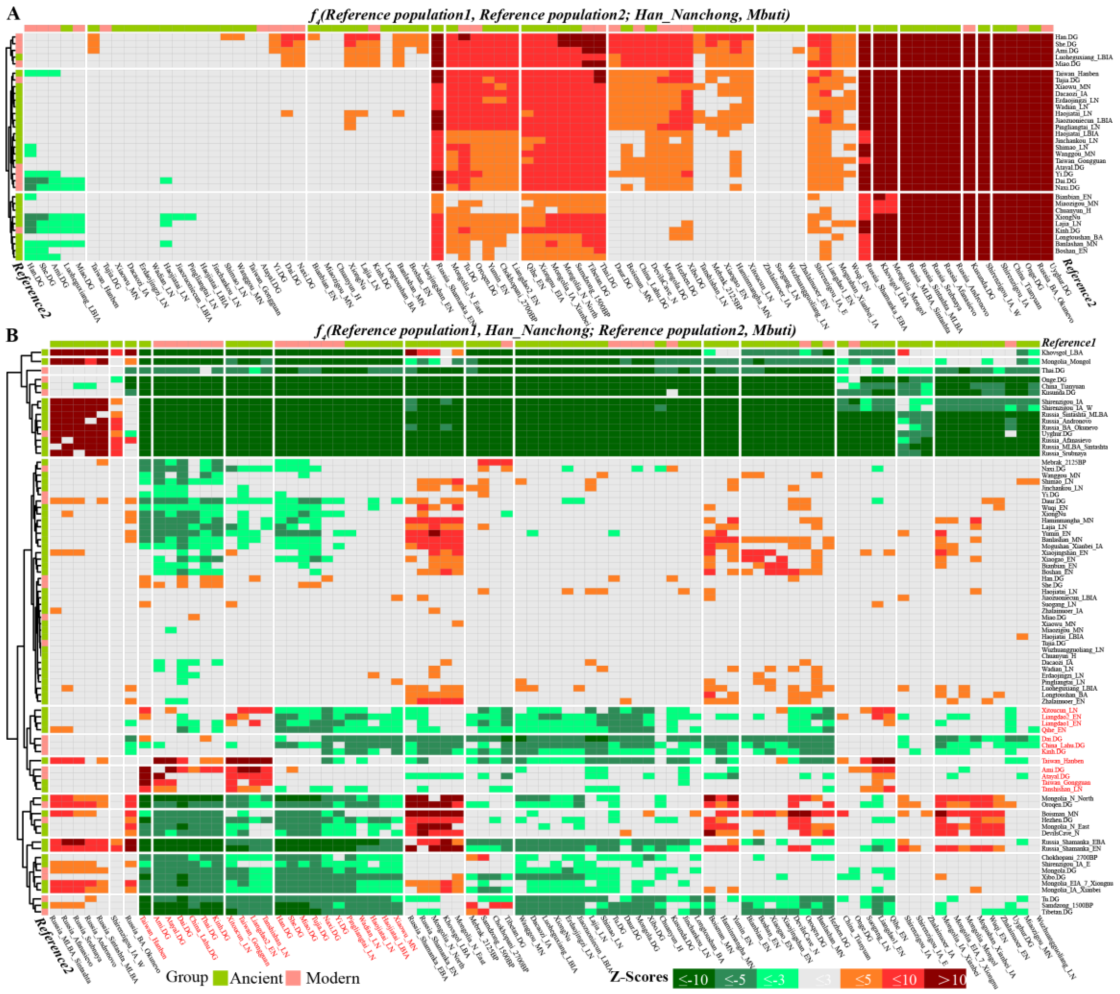


**Figure S7. Excess of sharing alleles between Nanchong Han and modern and ancient Eurasians.**

Symmetrical $f_4$-statistics in the form $f_4$*(Reference population1, reference population2; Han_Nanchong, Mbuti)* to test the genetic affinity between Eurasian reference populations (**A**). Affinity-$f_4$-statistics in the form $f_4$*(Reference population1, Han_Nanchong; reference population2, Mbuti)* to explore the genetic continuity and admixture between potential ancestral sources and Boshu Hui (**B**). The red color denoted the third population shared more derived alleles with the first population compared with the second one. And the green color denoted the third population shared more alleles with the second one relative to the first one.

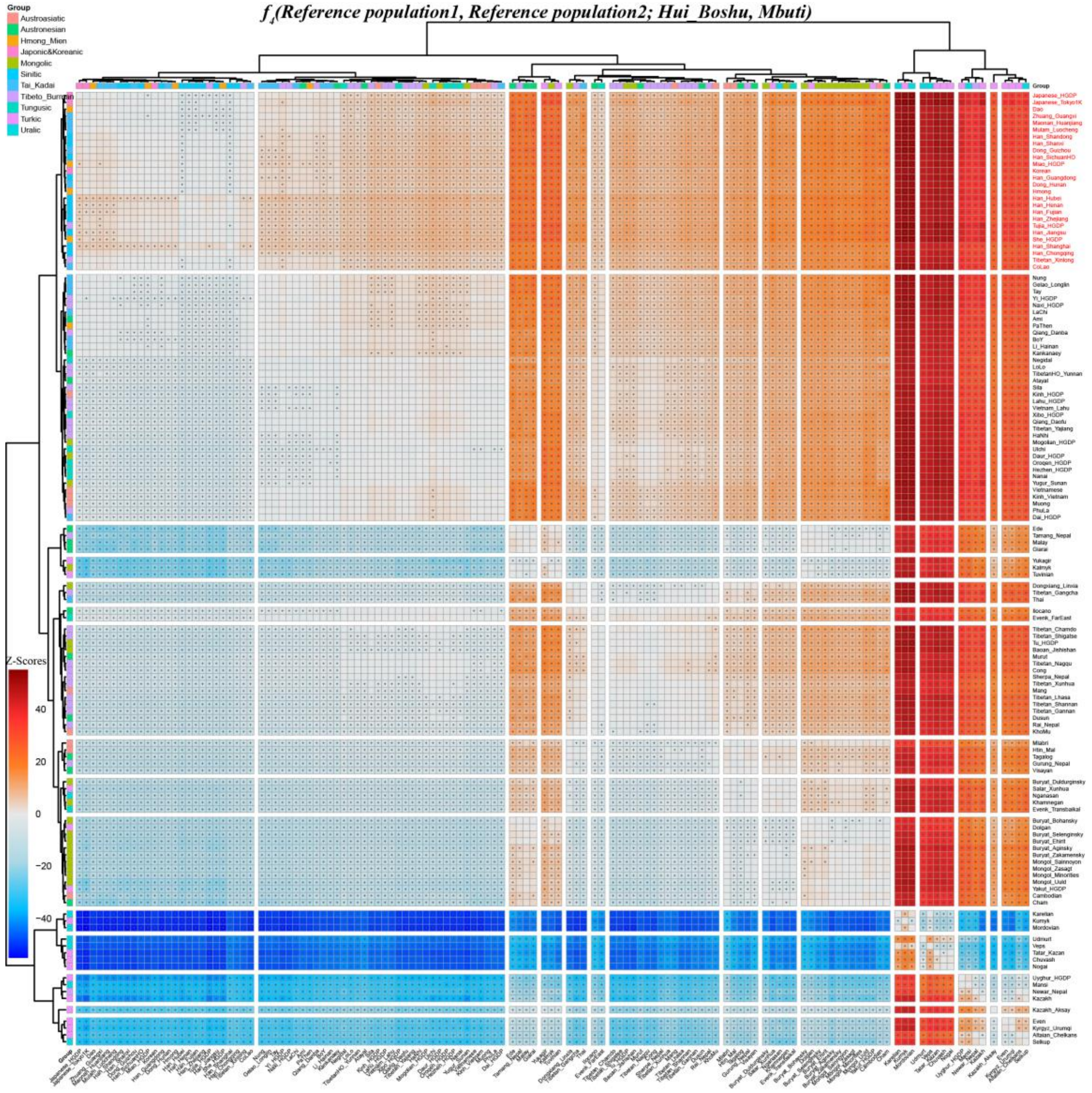


**Figure S8. Results of $f_4$-statistics showed genomic relationship inferred from *$f_4$(Reference population1, reference population2; Hui_Boshu, Mbuti)* based on the merged Human Origin dataset.**

Symmetrical $f_4$-statistics in the form *$f_4$(Reference population1, reference population2; Hui_Boshu, Mbuti)* to test excess allele sharing between Boshu Hui and northern East northern East Asians (Right population lists) relative to other Eurasian reference populations. Red color showed positive $f_4$ values, which suggested that Boshu Hui possessed significantly more allele sharing of reference popultion1 (Right population lists) related to reference population2 (Bottom population lists). Blue color showed negative $f_4$ values, which suggested that Boshu Hui harbored excess sharing alleles with reference population2 (Bottom population lists) relative to reference popultion1 (Right population lists). Statistically significant $f$-statistics were marked as "+".

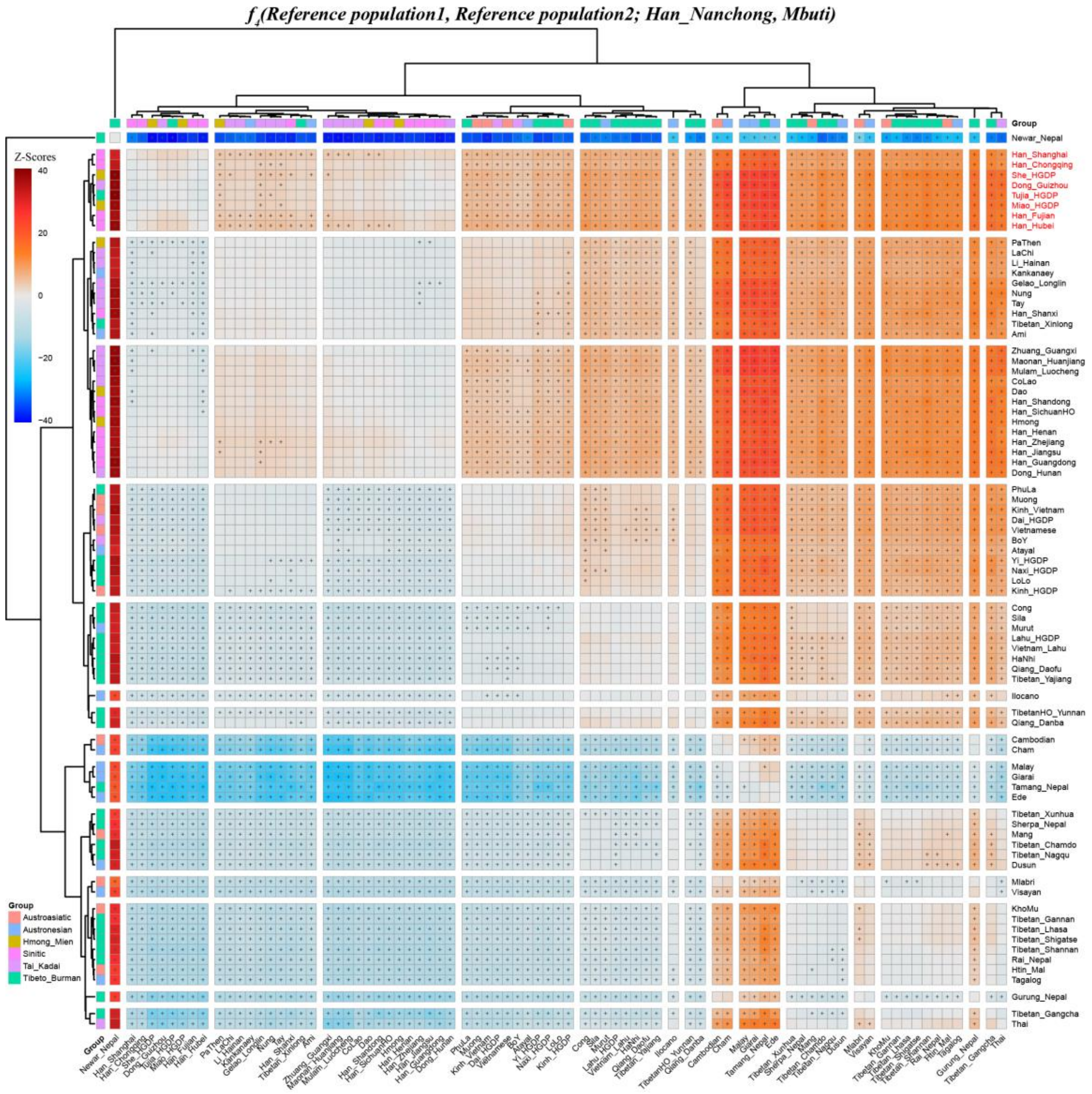


**Figure S9. Results of $f_4$-statistics showed genomic relationship inferred from *$f_4$(Reference population1, reference population2; Han_Nanchong, Mbuti)* based on the merged Human Origin dataset.**

Symmetrical $f_4$-statistics in the form *$f_4$(Reference population1, reference population2; Han_Nanchong, Mbuti)* to test excess allele sharing between Nanchong Han and northern East Asians (Right population lists) relative to other Eurasian reference populations. Red color showed positive $f_4$ values, which suggested that Nanchong Han possessed significantly more allele sharing of reference popultion1 (Right population lists) related to reference population2 (Bottom population lists). Blue color showed negative $f_4$ values, which suggested that Nanchong Han harbored excess sharing alleles with reference population2 (Bottom population lists) relative to reference popultion1 (Right population lists). Statistically significant *f*-statistics were marked as “+”.

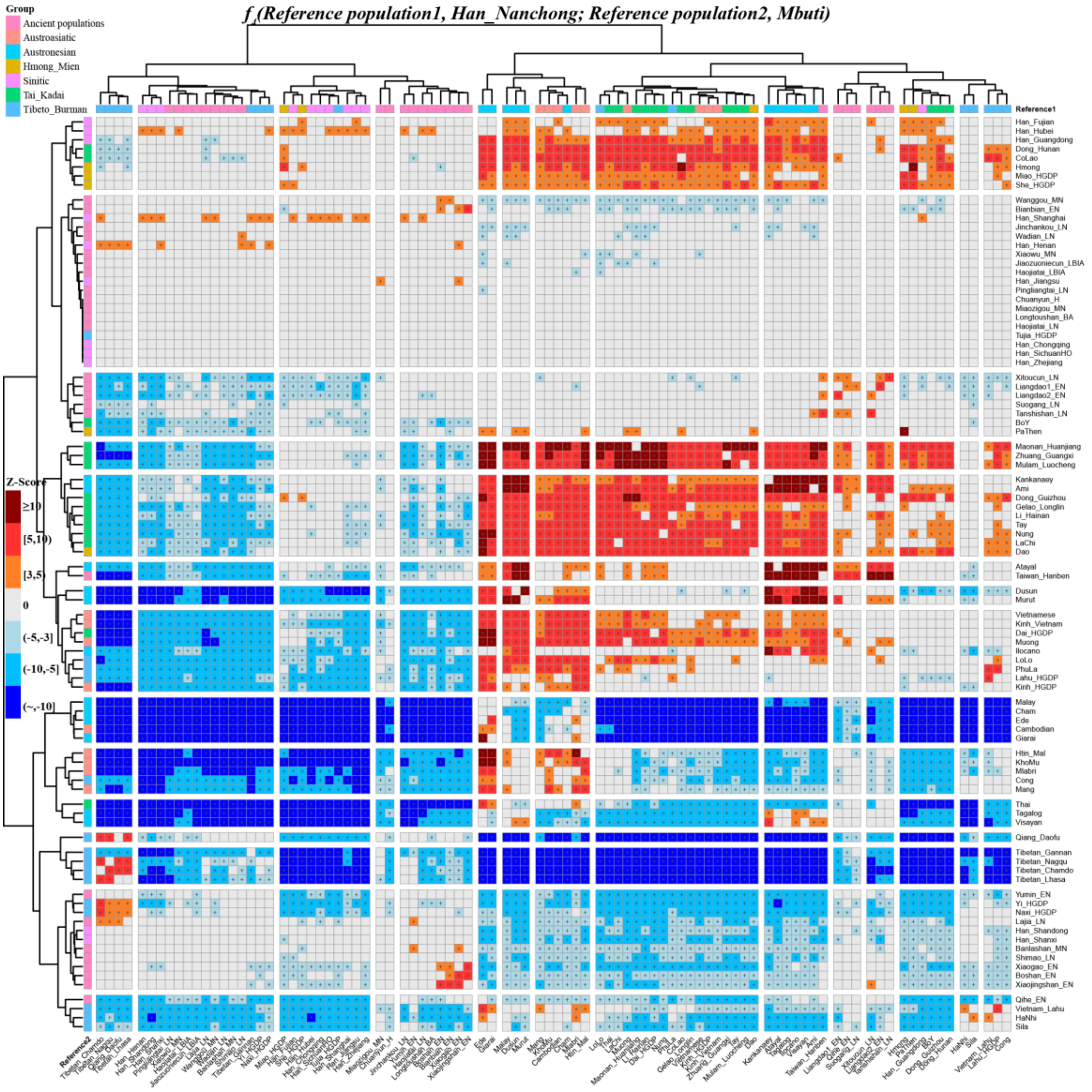


**Figure S10. Excess of sharing alleles between Nanchong Han and modern and ancient Eurasians based on the merged Human Origin dataset.**

Affinity-$f_4$-statistics in the form $f_4$*(Reference population1, Han_Nanchong; reference population2, Mbuti)* to explore the genetic continuity and admixture between potential ancestral sources and Nanchong Han based on the merged Human Origin dataset. The red color denoted the third population (Bottom population lists) shared more derived alleles with the first population (Right population lists) compared with the second one (Nanchong Han). And the green color denoted the third population shared more alleles with the Nanchong Han relative to the first one. Statistically significant *f*-statistics were marked as “+”.

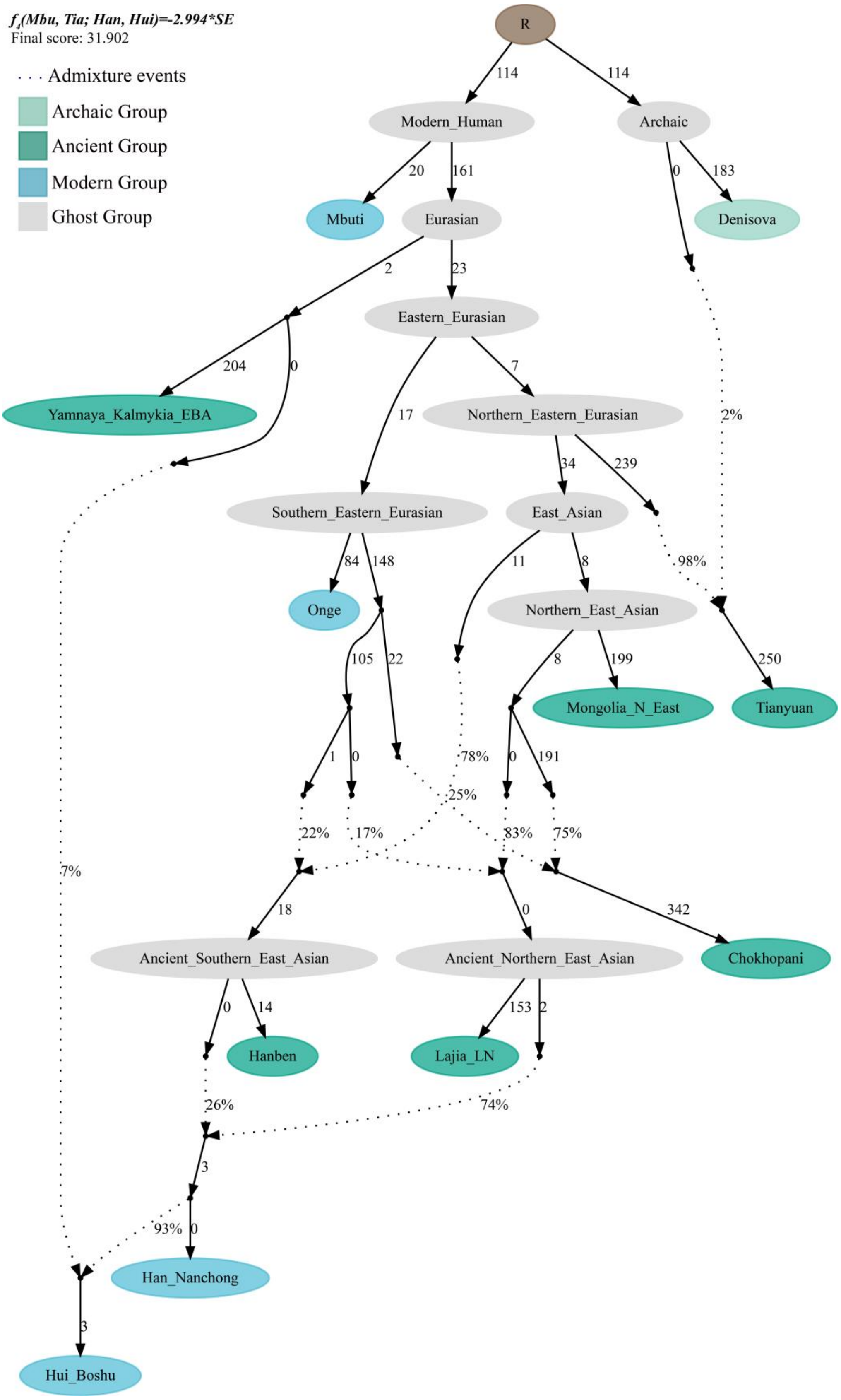


**Figure S11. *qpGraph*-based admixture graph showed the western Eurasian gene flow event related to Kalmykia Yamnaya into Boshu Hui.**

Branch length was marked with the $f_2$ shared drift distance (1000 times). Admixture events were denoted as dotted line. Admixture proportion was marked along the dotted line. EBA: Early Bronze Age; LN, Late Neolithic; N, Neolithic.

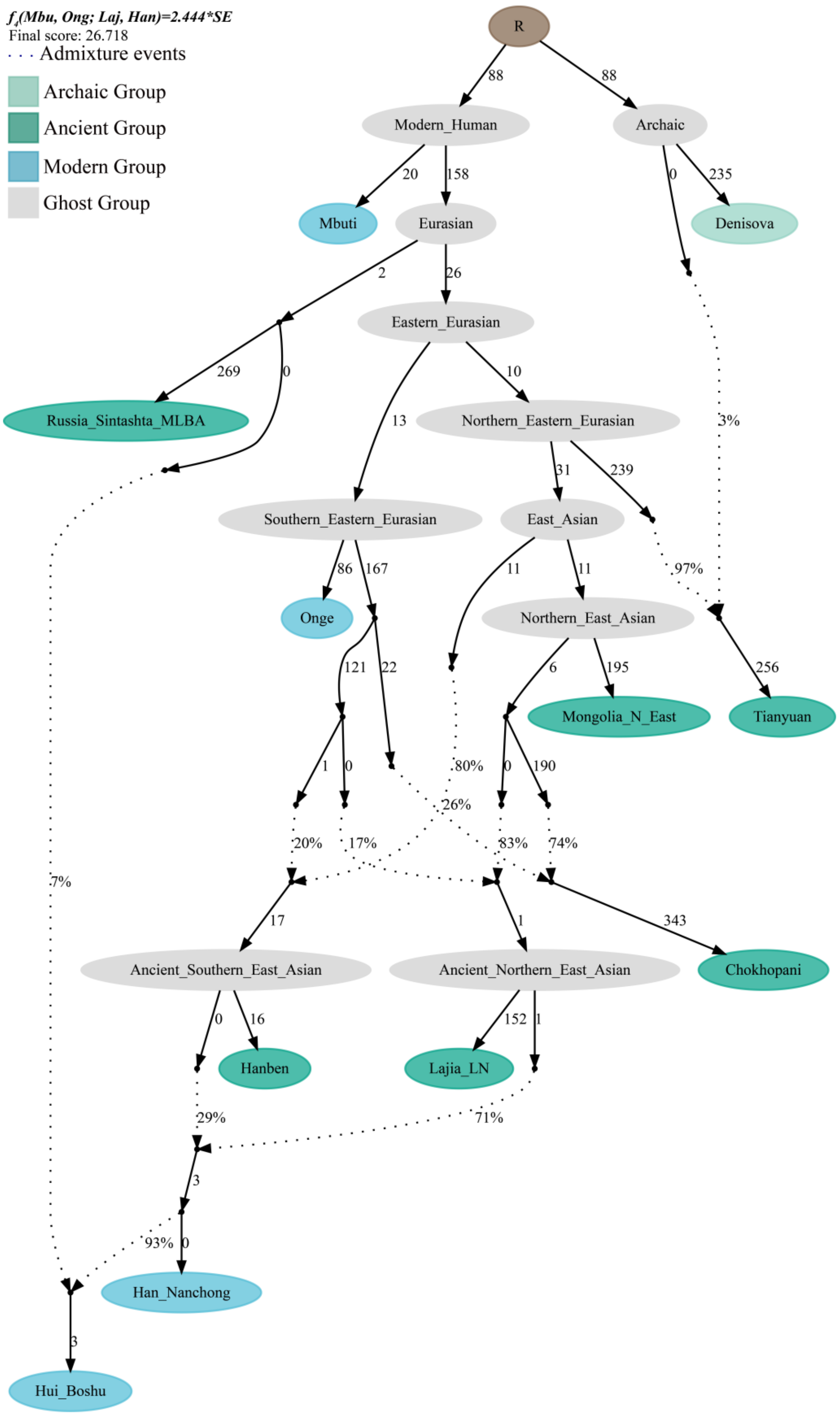


**Figure S12.** ***qpGraph*-based admixture graph showed the western Eurasian gene flow event related to Russia_Sintashta_MLBA into Boshu Hui.**

Branch length was marked with the $f_2$ shared drift distance (1000 times). Admixture events were denoted as dotted line. Admixture proportion was marked along the dotted line. MLBA: Middle-Late Bronze Age; LN, Late Neolithic; N, Neolithic.

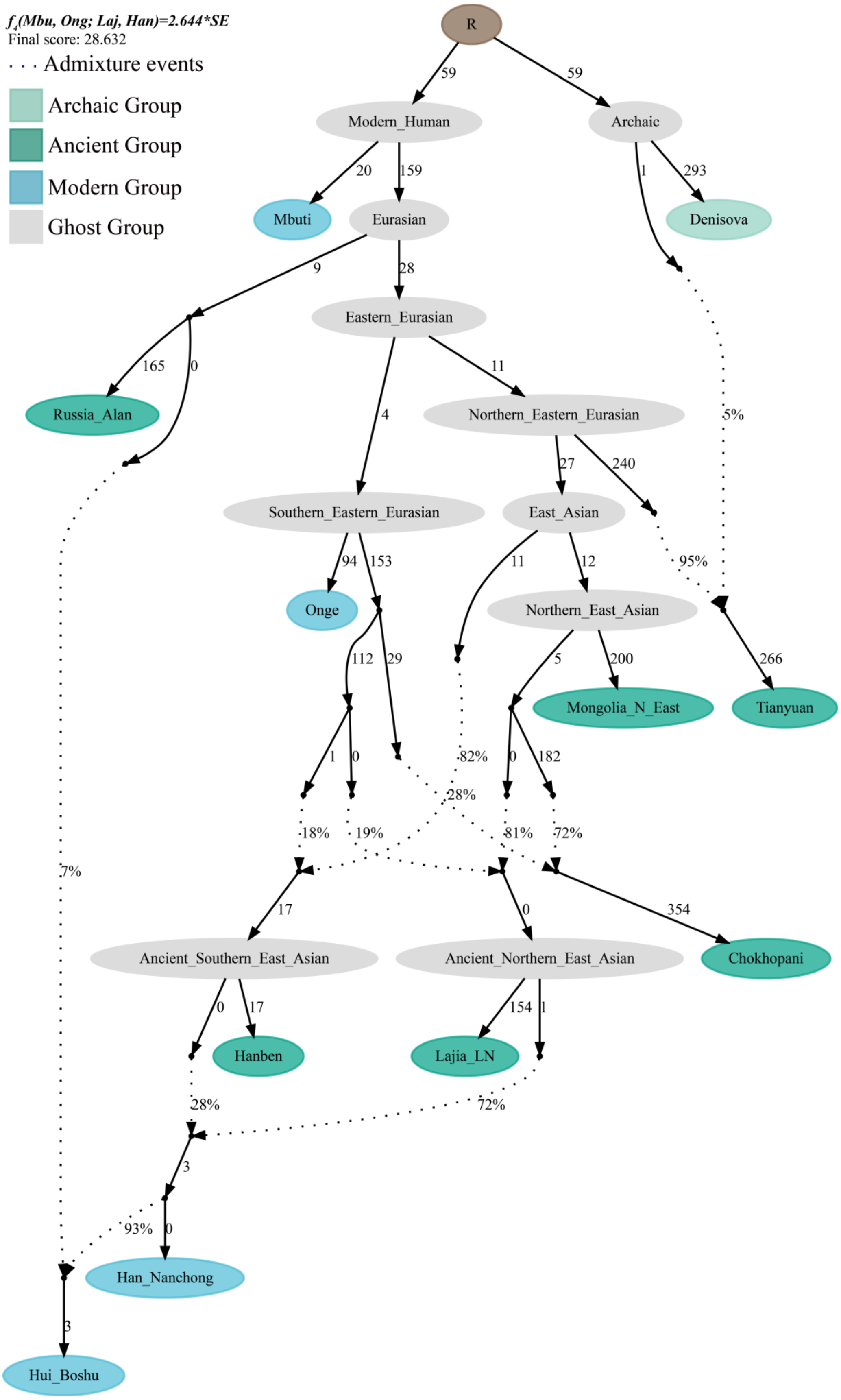


**Figure S13. *qpGraph*-based admixture graph showed the western Eurasian gene flow event related to Russia Alan into Boshu Hui.**

Branch length was marked with the $f_2$ shared drift distance (1000 times). Admixture events were denoted as dotted line. Admixture proportion was marked along the dotted line. LN, Late Neolithic; N, Neolithic.

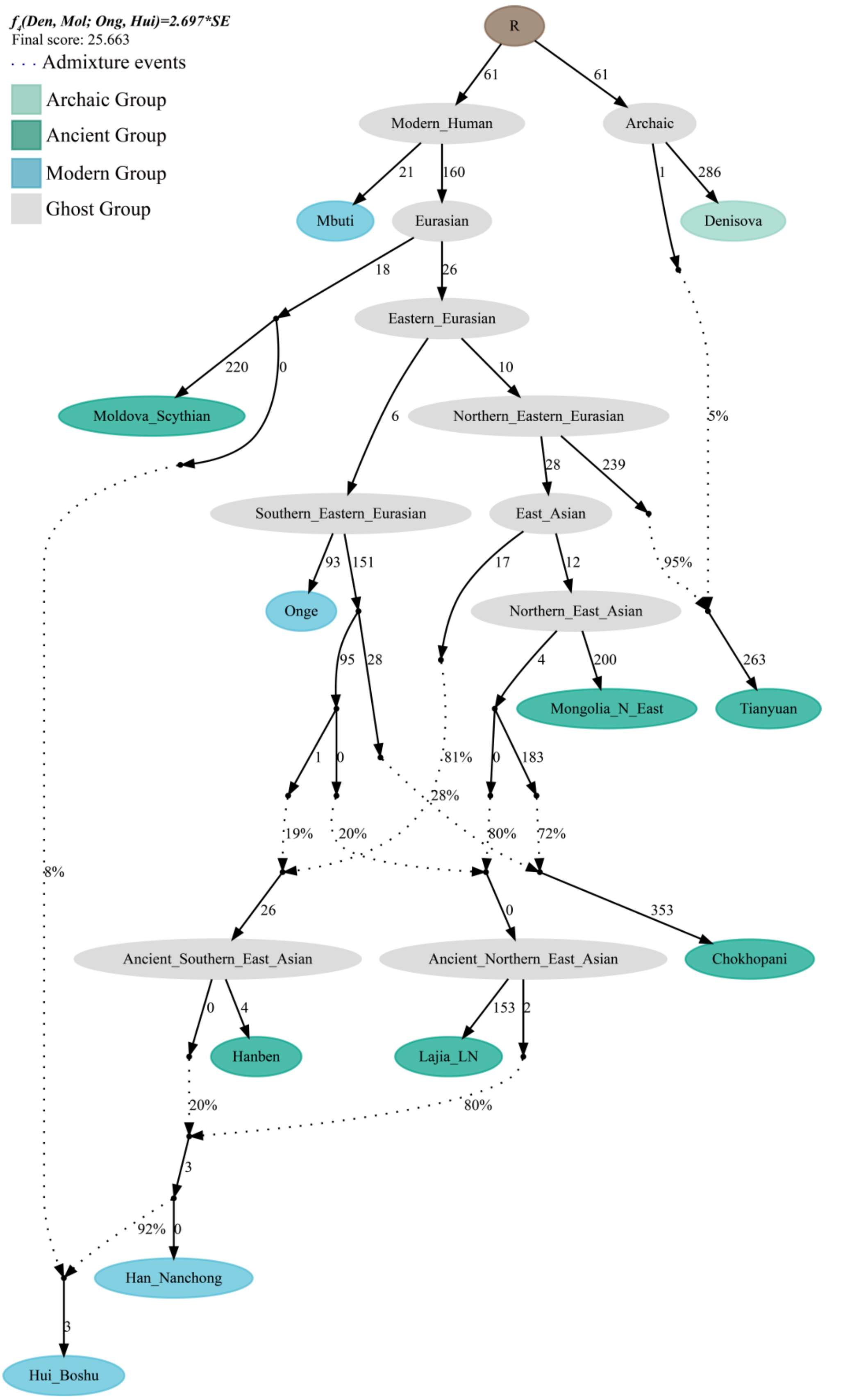


**Figure S14. *qpGraph*-based admixture graph showed the western Eurasian gene flow event related to Moldova Scythian into Boshu Hui.**

Branch length was marked with the $f_2$ shared drift distance (1000 times). Admixture events were denoted as dotted line. Admixture proportion was marked along the dotted line. LN, Late Neolithic; N, Neolithic.

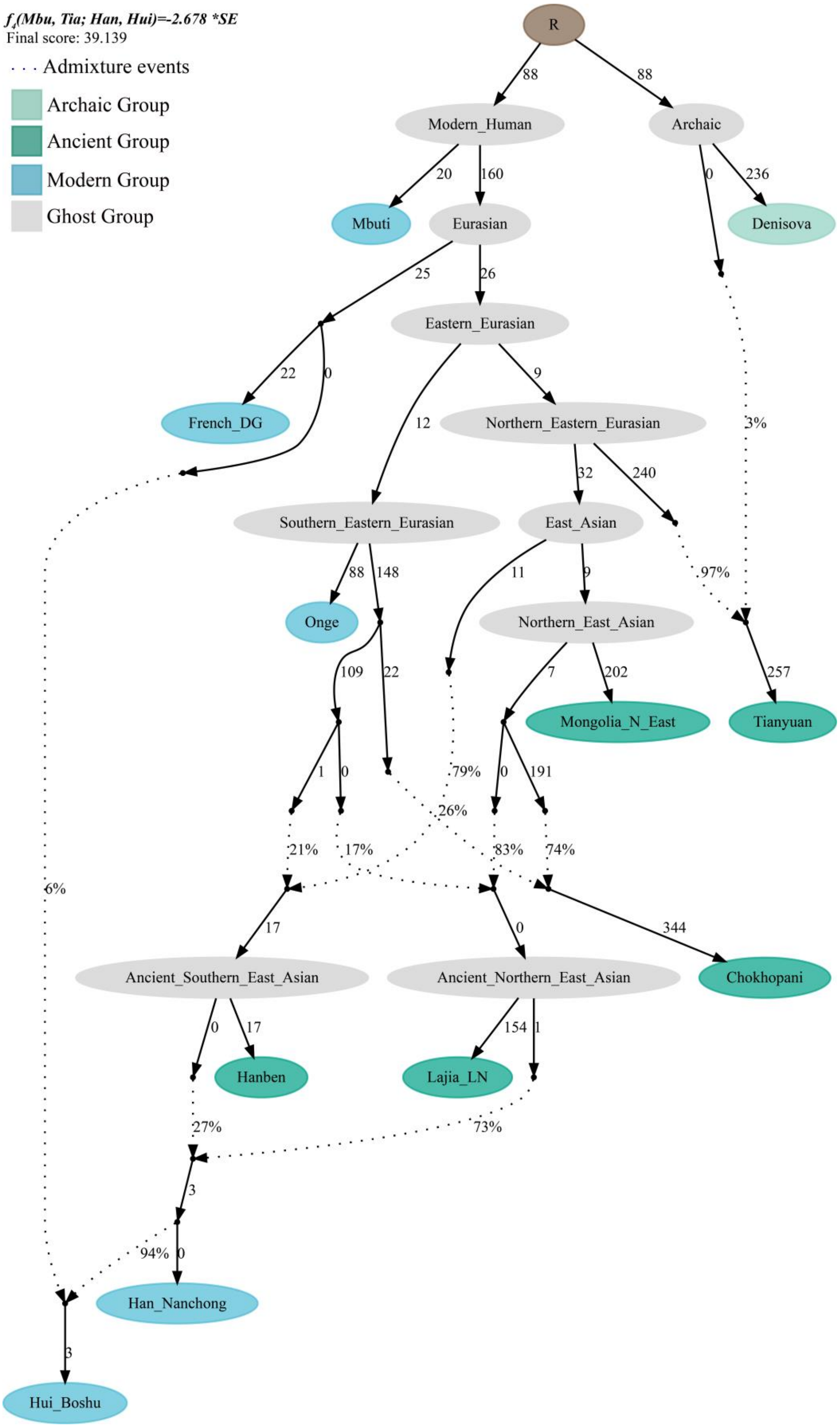


**Figure S15. *qpGraph*-based admixture graph showed the western Eurasian gene flow event related to French into Boshu Hui.**

Branch length was marked with the $f_2$ shared drift distance (1000 times). Admixture events were denoted as dotted line. Admixture proportion was marked along the dotted line. LN, Late Neolithic; N, Neolithic.

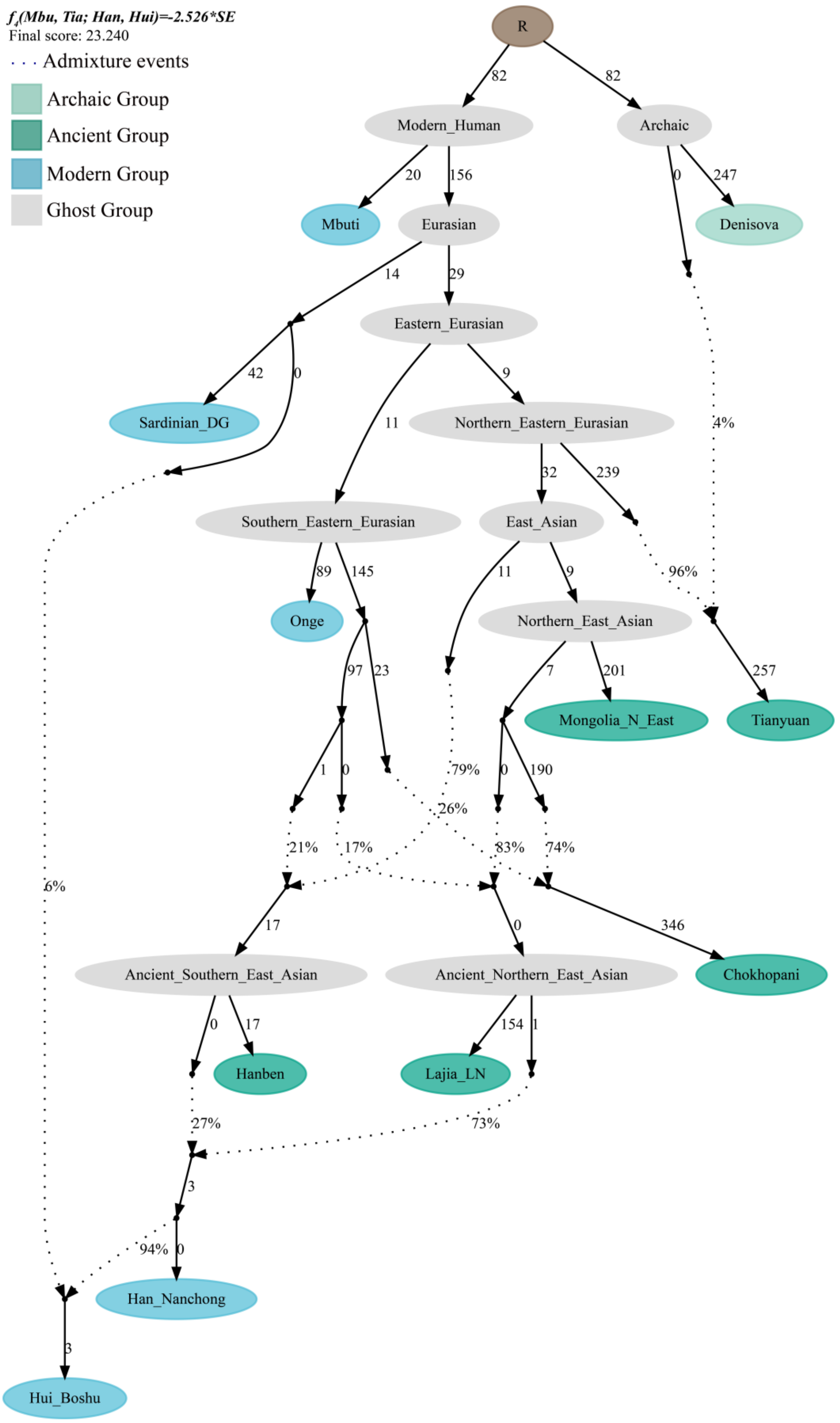


**Figure S16. *qpGraph*-based admixture graph showed the western Eurasian gene flow event related to Sardinian into Boshu Hui.**

Branch length was marked with the $f_2$ shared drift distance (1000 times). Admixture events were denoted as dotted line. Admixture proportion was marked along the dotted line. LN, Late Neolithic; N, Neolithic.

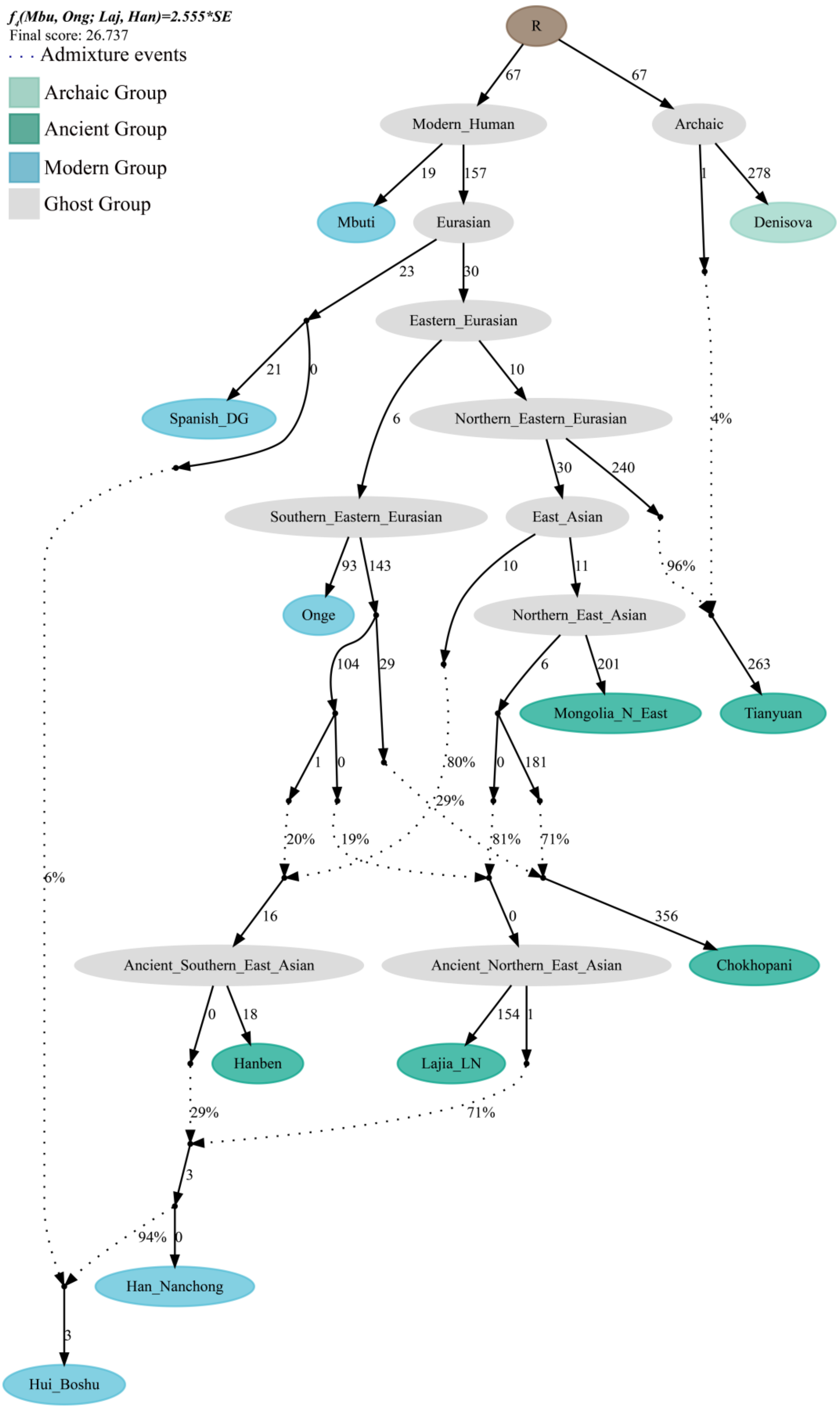


**Figure S17. *qpGraph*-based admixture graph showed the western Eurasian gene flow event related to Spanish into Boshu Hui.**

Branch length was marked with the $f_2$ shared drift distance (1000 times). Admixture events were denoted as dotted line. Admixture proportion was marked along the dotted line. LN, Late Neolithic; N, Neolithic.

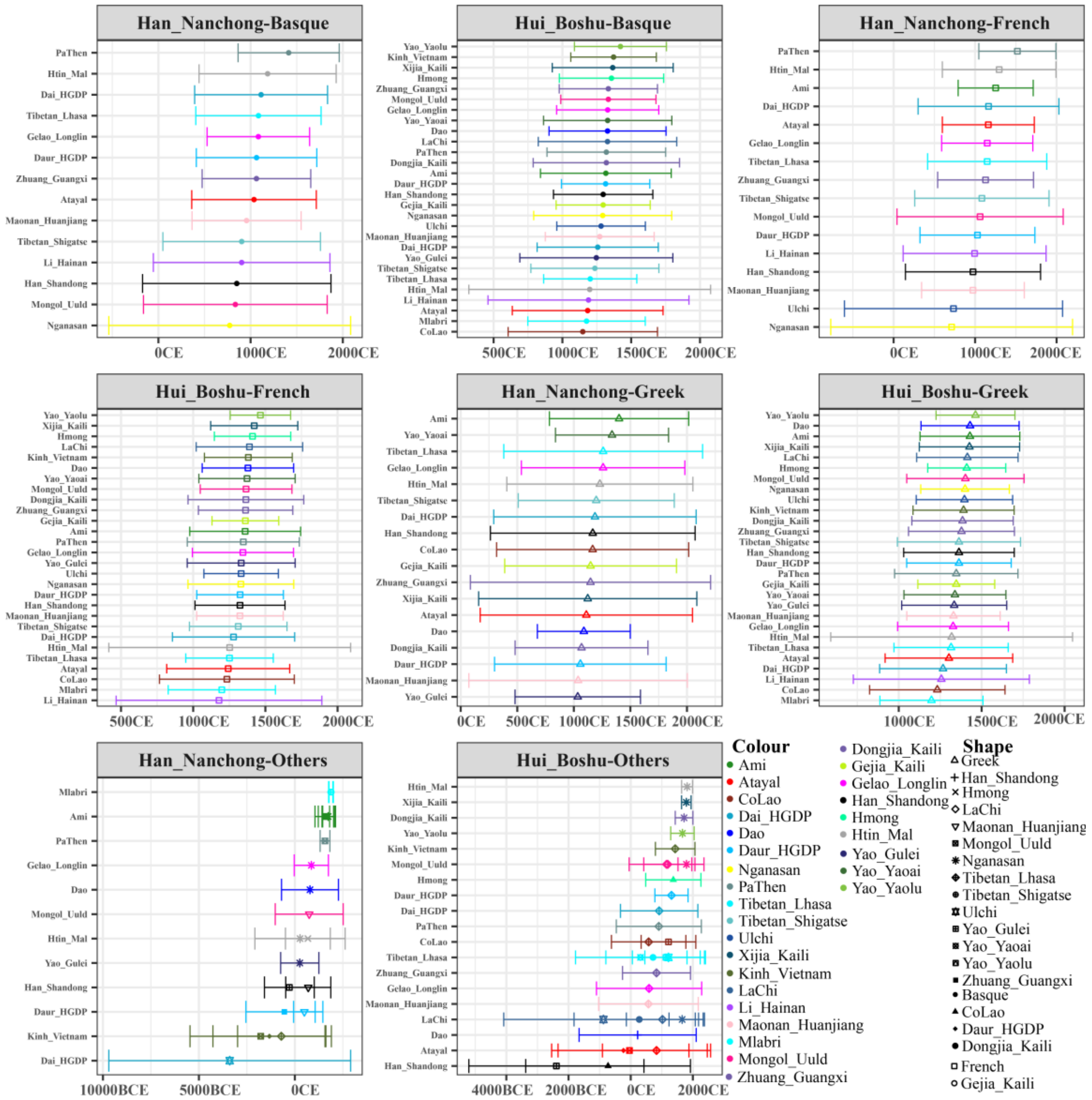


**Figure S18. Admixture-introduced linkage disequilibrium (ALDER) based admixture time between different northern and southern or eastern and western ancestral sources.**

We used 28 years as the one generation length. All marked years in the bottom was calculated using the formula as Year=1950-28*(Generation-1). All comprehensive raw data were presented in Supplementary **Table S17**
